# Distribution of ACE2, CD147, cyclophilins, CD26 and other SARS-CoV-2 associated molecules in human tissues and immune cells in health and disease

**DOI:** 10.1101/2020.05.14.090332

**Authors:** U. Radzikowska, M. Ding, G. Tan, D. Zhakparov, Y. Peng, P. Wawrzyniak, M. Wang, S. Li, H. Morita, C. Altunbulakli, M. Reiger, AU. Neumann, N. Lunjani, C. Traidl-Hoffmann, K. Nadeau, L. O’Mahony, CA. Akdis, M. Sokolowska

**Affiliations:** Swiss Institute of Allergy and Asthma Research (SIAF), University of Zurich, Davos, Switzerland; Christine Kühne – Center for Research and Education (CK-CARE), Davos, Switzerland; Department of Regenerative Medicine and Immune Regulation, Medical University of Bialystok, Bialystok, Poland; Department of Allergology, Zhongnan Hospital of Wuhan University, Wuhan, China; Functional Genomic Centre Zurich, ETH Zurich/University of Zurich, Zurich, Switzerland; Otorhinolaryngology Hospital, The First Affiliated Hospital, Sun Yat-sen University, Guangzhou, China; Division of Clinical Chemistry and Biochemistry, University Children’s Hospital Zurich, Zurich, Switzerland; Children’s Research Center, University Children’s Hospital Zurich, Zurich, Switzerland; Department of Otolaryngology, Head and Neck Surgery, Beijing TongRen Hospital, Capital Medical University and the Beijing Key Laboratory of Nasal Diseases, Beijing Institute of Otolaryngology, Beijing, China; Department of Cancer Immunology, Institute for Cancer Research, Oslo University Hospital, Oslo, Norway; Department of Allergy and Clinical Immunology, National Research Institute for Child Health and Development, Tokyo, Japan; Chair and Institute of Environmental Medicine, UNIKA-T, Technical University of Munich and Helmholtz Zentrum Munchen, Augsburg, Germany; Institute of Computational Biology (ICB), Helmholtz Zentrum Munchen, Munich, Germany; Institute of Experimental Medicine (IEM), Czech Academy of Sciences, Prague, Czech Republic; Sean N Parker Centre for Allergy and Asthma Research at Stanford University, Department of Medicine, Stanford University School of Medicine, Stanford, USA; Department of Medicine and School of Microbiology, APC Microbiome Ireland, National University of Ireland, Cork, Ireland

**Author notes:** equal contribution (U. Radzikowska, M. Ding). **corresponding author:** Milena Sokolowska MD, PhD, Head Immune Metabolism, Swiss Institute of Allergy and Asthma Research (SIAF), University of Zurich, Herman-Burchard-Strasse 9, CH-7265 Davos-Wolfgang, https://www.siaf.uzh.ch/immune_metabolism.html.

**Keywords:** COVID-19, SARS receptor, obesity, hypertension, asthma, COPD, comorbidity

## Abstract

**Background:** Morbidity and mortality from COVID-19 caused by novel coronavirus SARS-CoV-2 is accelerating worldwide and novel clinical presentations of COVID-19 are often reported. The range of human cells and tissues targeted by SARS-CoV-2, its potential receptors and associated regulating factors are still largely unknown. The aim of our study was to analyze the expression of known and potential SARS-CoV-2 receptors and related molecules in the extensive collection of primary human cells and tissues from healthy subjects of different age and from patients with risk factors and known comorbidities of COVID-19.

**Methods:** We performed RNA sequencing and explored available RNA-Seq databases to study gene expression and co-expression of ACE2, CD147 (*BSG*), CD26 (*DPP4*) and their direct and indirect molecular partners in primary human bronchial epithelial cells, bronchial and skin biopsies, bronchoalveolar lavage fluid, whole blood, peripheral blood mononuclear cells (PBMCs), monocytes, neutrophils, DCs, NK cells, ILC1, ILC2, ILC3, CD4^+^ and CD8^+^ T cells, B cells and plasmablasts. We analyzed the material from healthy children and adults, and from adults in relation to their disease or COVID-19 risk factor status.

**Results:** *ACE2* and *TMPRSS2* were coexpressed at the epithelial sites of the lung and skin, whereas CD147 (*BSG*), cyclophilins (*PPIA and PPIB*), CD26 (*DPP4*) and related molecules were expressed in both, epithelium and in immune cells. We also observed a distinct age-related expression profile of these genes in the PBMCs and T cells from healthy children and adults. Asthma, COPD, hypertension, smoking, obesity, and male gender status generally led to the higher expression of ACE2- and CD147-related genes in the bronchial biopsy, BAL or blood. Additionally, CD147-related genes correlated positively with age and BMI. Interestingly, we also observed higher expression of ACE2- and CD147-related genes in the lesional skin of patients with atopic dermatitis.

**Conclusions:** Our data suggest different receptor repertoire potentially involved in the SARS-CoV-2 infection at the epithelial barriers and in the immune cells. Altered expression of these receptors related with age, gender, obesity and smoking, as well as with the disease status might contribute to COVID-19 morbidity and severity patterns.

## Introduction

A novel coronavirus SARS-CoV-2 leading to COVID-19 was identified for the first time in December 2019 and as of today has infected already more than 4.3 million and killed more than 290 000 people worldwide (as of 13^th^ of May 2020)^1^. SARS-CoV-2 virus has very high genome sequence similarity to two other human coronaviruses SARS-CoV and MERS-CoV^2^. Therefore, it is highly possible that SARS-CoV-2 uses similar approaches to cell entry and replication in various cells and tissues. In fact, it is already known that SARS-CoV-2 uses the same receptor ACE2 to enter the cells via its structural spike glycoprotein (S), yet with much higher affinity, which might translate to the massive SARS-CoV-2 spread as compared to SARS-CoV^3^. Host transmembrane protease serine 2 (TMPRSS2; encoded by *TMPRSS2*) cleaves spike protein into two subunits, which is a necessary step for the virus fusion to cellular membranes and entry the cell^3^. Recent structural studies revealed that this process can be potentially inhibited by B0AT1 (S6A19; encoded by *SLC6A19*), an amino acid transporter, presence of which may block the access of TMPRSS2 to the cleavage site on ACE2^4^ (Table S1).

The expression of ACE2 (encoded by *ACE2)* is very ubiquitous in the lung, heart, kidney and intestine, but it is rarely expressed in immune cells^5,6^. However, immune cells can be potentially infected by SARS-CoV-2, as in case of MERS-CoV and SARS-CoV^7,8^. Thus, it is highly possible that there are other receptors for virus entry in different cell types. Indeed, similarly with what has been shown in SARS-CoV, another receptor - CD147, called also basigin (encoded by *BSG*), has been recently shown to act as a receptor for SARS-CoV-2 in T cell lines and in cell lines of epithelial origin^9,10^. Interestingly, the same receptor is a putative receptor not only for SARS-CoV, but also for HIV-1 and measles, as well as is a receptor for malaria entry to erythrocytes^11–14^. An anti-CD147 antibody that blocks infection with SARS-CoV-2 *in vitro* and its humanized drug (meplazumab) has been already used in a clinical trial in patients with COVID-19 pneumonia. Meplazumab seemed to facilitate viral clearance, return to normal levels of lymphocyte count and decrease in CRP^9,15^. The percentage of improvement in patients with severe and critical COVID-19 presentations seemed higher in weekly meplazumab treatment compared to patients on the conventional treatment^15^.

CD147 is a transmembrane receptor interacting with several extracellular and intracellular partners forming a transmembrane supramolecular complex^16^. A group of extracellular molecules can bind and activate CD147 are cyclophilins A and B (encoded by *PPIA* and *PPIB*), acute phase protein S100A9, E-selectin (encoded by *SELE)* and platelet glycoprotein VI (encoded by *GP6*)^17–20^. Additionally, CD147 has three Asn glycosylation sites, to which high mannose-type and complex-type glycans might bind^21^. Spike protein of SARS-CoV-2 is highly glycosylated^22^, which increases the chance of binding to the cells. It is possible that the virus can only attach to the cell surface and induce different cellular programs, leading to cell overactivation, exhaustion and death, especially in the severe phase of the disease, where cytokine storm and lymphopenia is commonly reported^23^.

Cyclophilins A and B have been shown to interact with non-structural protein 1 (nsp1) of SARS-CoV intracellularly^24,25^. They can be incorporated to the viral capsid and released, which further enables virus binding to CD147 and subsequent infection of CD147-expressing cells. Cyclophilins play a critical role in the replication process of HIV-1, HCV and many other viruses^12,26^. Cyclosporine A, a strong immunosuppressive agent acts mainly via binding to cellular cyclophilins. In the same mechanism, cyclosporine A suppresses replication of various coronaviruses^27^. Thus, cyclophilins were identified as targets for pan-coronavirus inhibitors^28^. Nevertheless, it is still not known if cyclophilins can interact with SARS-CoV-2.

Intracellular and transmembrane partners of CD147 are equally important in the infection process of HIV, measles and SARS-CoV. It has been shown that truncation of cytoplasmic tail of CD147 prevents HIV infection^29^. Therefore, it is possible that these molecules can take part and regulate the process of SARS-CoV-2 entry through CD147 in a comparable way as TMPRSS2 and SLC6A19 for ACE2. Main transmembrane partners of CD147 with direct binding sites to CD147 are monocarboxylate transporters 1-4 (MCT1-MCT4; encoded by: *SLC16A1, SLC16A7, SLC16A8, SLC16A3*), main glucose transporter, GLUT-1 (encoded by *SLC2A1)* and CD44 (encoded by *CD44*), an important receptor for hyaluronan-main component of extracellular matrix^30–32^. CD147 also interacts intracellularly with integrins α3β1 and α6β1 (encoded by *ITGA3, ITGA6, ITGB1)* indirectly via CD98 (*SLC7A5*), CD43 (*SPN*), MCT4 and galectin-3 (*LGALS3*)^33–36^. In addition, CD147 suppresses two important protein complexes by direct interaction: NOD2 (*NOD2*), an important innate immunity component and gamma-secretase complex, responsible for cleavage of beta-amyloid precursor from plasma membranes, encoded by *PSEN1, NCSTN, APH1A, APH1B, PSENEN*^37,38^. Due to the interactions with so many molecules, CD147 plays a crucial role in energy metabolism of the cell, motility, recruitment and activation, although the final outcome of its stimulation depends on the cell type and co-expression of the other molecules. However, the co-expression of these molecules in different cell types and across various human tissues is not known.

Importantly, in T cells CD147 suppresses T-cell receptor-dependent activation mainly by inhibition of nuclear factor of activated T-cells (NFATs)^39,40^. The NFAT family consists of five proteins: NF-ATc1 (encoded by *NFATC1*), NF-ATc2 (encoded by *NFATC2*), NF-ATc3 (encoded by *NFATC3*), NF-ATc4 (encoded by *NFATC4*) and NF-AT5 (encoded by *NFAT5*)^41,42^. NFAT complex is a major transcriptional regulator in naive T cells and differentiated effector T cells, dependent on calcium/PLCγ/calmodulin/calcineurin signaling^41,42^. It is also crucial in regulation of T cell anergy and in differentiation and function of T regulatory (Treg) cells^42^. CD147 has been reported as a marker of human Treg cells with highly suppressive activity^43^. NFAT signaling is also important in other cell types such as DCs, mast cells and B cells^41^. Since cyclophilins are regulators of NFAT activation, the extracellular binding of the virus through cyclophilin-CD147 complex, as well as intracellular interactions of cyclophilins with the virus proteins might be important in COVID-19 development.

CD26 (encoded by *DPP4*), has emerged recently as a potential receptor for SARS-CoV-2, due to the fact that it is a main cellular entry for MERS-CoV^44,45^. Recent structural studies predict that SARS-CoV-2 spike protein directly interact with CD26 on the host cells^22^. CD26 is a cell surface glycoprotein involved in T-cell receptor-mediated T-cell activation and proliferation. It is highly expressed in CD4 and CD8 T cells, and in lower quantities also in NK cells and DCs. As an additional function, it acts as a serine exopeptidase, cleaving peptides of various chemokines, growth factors and peptide hormones. Interestingly, it is also involved in extracellular matrix cleavage. It has been shown in human cells that several polymorphisms in CD26 gene reduce entry of MERS into the host cells^46^. It remains to be functionally determined if this protein can be a functional receptor for SARS-CoV-2.

The aim of our study was to analyze the gene expression of ACE2, CD147, cyclophilins, CD26 and other SARS-CoV-2-related molecules in the broad range of primary human innate and adaptive immune cells and tissues, based on our own next generation sequencing data and public databases from different human cell types and across the diseases which are known to predispose to COVID-19.

## Material and Methods

### Study subjects, samples and study description

We analyzed gene expression of SARS-CoV-2 receptors and related molecules’ (Table S1) in a broad range of tissues and immune cells from the human RNA-seq databases generated by our *ex vivo* and *in vitro* approaches in the Swiss Institute of Asthma and Allergy Research (SIAF), by our collaborators or from the Gene Expression Omnibus (http://www.ncbi.nlm.nih.gov/). Baseline gene expression was evaluated *in vitro* in air liquid interface (ALI) - differentiated human primary bronchial epithelial cells (HBECs) (n=10). Direct *ex vivo* analyses of non-diseased human tissues included bronchial biopsies (n=5), bronchoalveolar lavage (BAL) cells (n=5) and skin biopsies (n=6). The other *ex vivo* investigated samples included whole blood (n=5), neutrophils (n=4), classical monocytes (n=4), plasmacytoid dendritic cells (pDCs) (n=4), innate lymphoid cells (ILC) 1, 2 and 3 (n=6 for each), natural killer cells (NK) (n=4), naïve CD4+ T cells (n=4), terminal effector CD4+ T cells (n=2), naïve CD8+ T cells (n=4), effector memory CD8+ T cells (n=4), naïve B cells (n=4) and plasmablasts (n=4) (for sorting markers please see Table S2).

We further investigated different expression patterns of ACE-2-, CD147- and CD26-related genes in the context of potential COVID-19 risk factors, namely age, gender, smoking status, diagnosis of asthma, chronic obstructive pulmonary disease (COPD), hypertension, obesity, atopic dermatitis (AD) (Table S3). Briefly, we performed in-depth curated analysis of HBECs (Control = 5, Asthma = 6, COPD =5), bronchial biopsies (Control = 16, Asthma = 22, COPD = 3, Non-obese = 20, Obese = 21, Normotension = 32, Hypertension = 9, Non-smoker = 19, Smoker = 21, Female = 14, Mal e= 27), BAL fluid (Control = 16, Asthma = 22, COPD = 2, Non-obese = 19 Obese = 21, Normotension = 31, Hypertension = 9, Non-smoker = 19, Smoker = 20, Female = 14, Male = 26), whole blood (Control = 17, Asthma = 21, COPD = 3, Non-obese = 20, Obese = 21, Normotension = 32, Hypertension = 9, Non-smoker = 19, Smoker = 21, Female = 14, Male = 27). In addition, we analyzed gene expression in PBMCs from infants and young children at age of 5-17 or 12-36 months (n=21 and 14, respectively), older children and adolescents at age of 4-16 years (n=16) and adults at age of 16-67 years (n=19), naïve CD4^+^ T cells from children at age 12 months (n=18) and adults at age 20-35 years (n=4) and skin biopsies (control: n=6, atopic dermatitis: non lesional sites n=11, lesional sites n=11). All studies were accompanied by the relevant ethical permissions, given by the appropriate Institutional Review Board. Each control and diseased subject gave informed consent. Total RNA from evaluated cells and tissues was extracted and transcriptome was analyzed with use of RNA-seq approaches.

### Data processing

Signatures of ACE2-, CD147- and CD26-related genes were curated from GSEA and MSigDB Database (Broad Institute, Massachusetts Institute of Technology, and Regents of the University of California) and from literature. Full sets of analyzed genes are described in the Table S1. Genes of interest were extracted from all data sets. RNA-seq data were processed with inhouse workflow available at https://github.com/uzh/ezRun. Significance threshold for differentially expressed genes was set to p < 0.05 and was calculated for the entire gene lists in each project. All calculations between different conditions were done using the edgeR R package^47^. Spearman correlation coefficient was calculated using Hmisc R package, with the threshold for significance set to α = 0.05. Correlation plots were done using Python’s Seaborn library. Co-expression heatmaps as well as correlation heatmaps were done using the corrplot R package

Detailed description of each study is included in the Supplementary Methods and the Table S3.

## Results

### ACE2 and related molecules are exclusively expressed at the barrier sites, whereas CD147, cyclophilins, CD26 and their interaction partners are ubiquitously expressed in immune cells and in epithelium

First, we analyzed the baseline expression and co-expression of ACE2-, CD147- and CD26-related genes in several immune cell types from control, non-diseased adult individuals. In agreement with other recent reports, we observed the expression of *ACE2* mainly in epithelial tissues, such as human bronchial epithelial cells and bronchial and skin biopsies (Figure 1A and S1)^6,48^. *ACE2* was more abundant in the lung, when compared with the skin barrier sides. In bronchial biopsies *ACE2* was co-expressed with *TMPRSS2* (Figure S2). Also, in the skin *TMPRSS2* was very prominent, while we did not find *SLC6A19* in any of the analyzed cells and tissues (Figure 1B and S1).

None of the analyzed immune cells expressed *ACE2* (Figure 1A), *TMPRSS2* (Figure 1B) or *SLC6A19* (Figures S1-3), either in the local lung environment (BAL cells) or in the blood.

**Figure 1.**
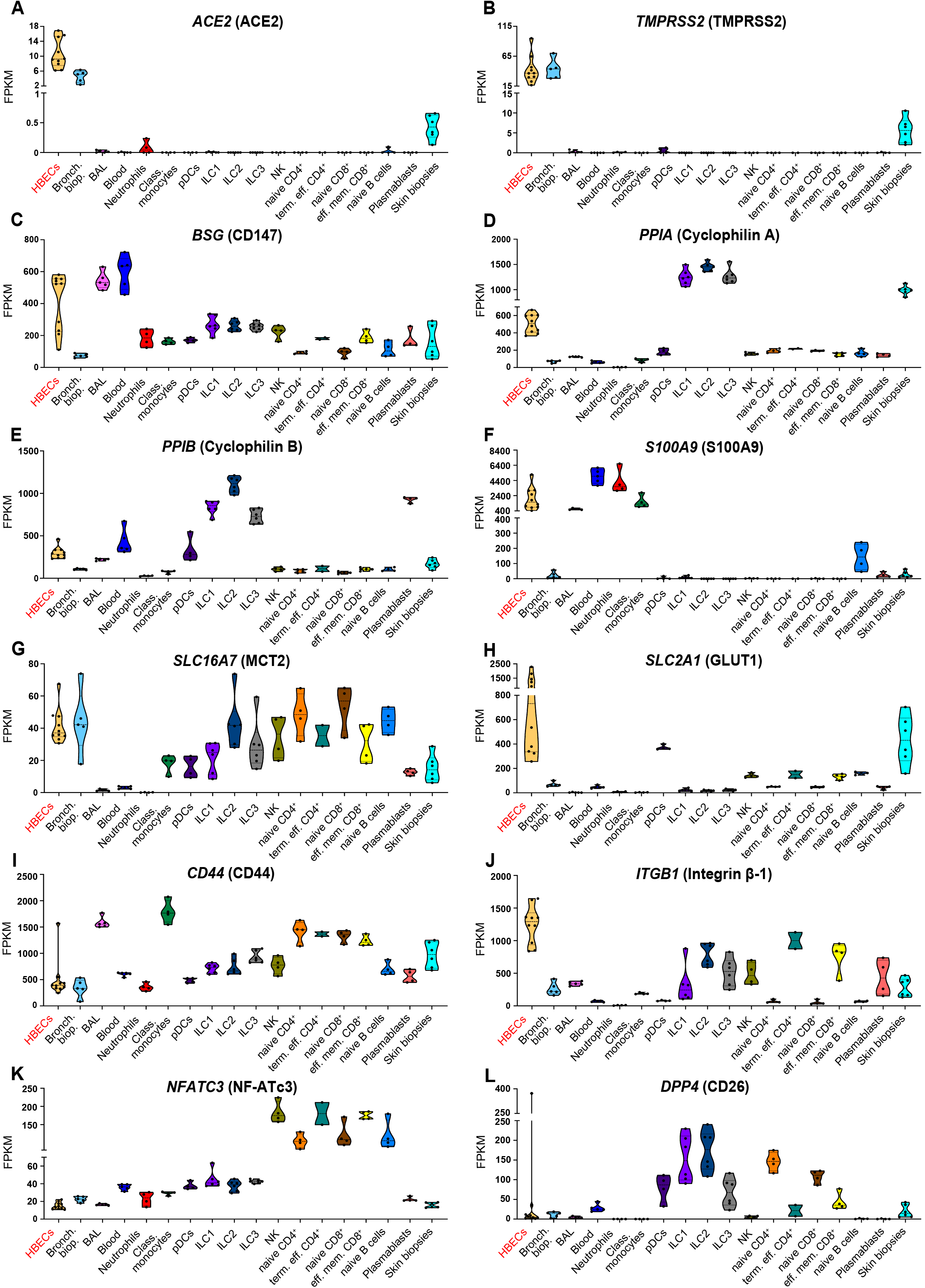
ACE2 and TMPRSS2 are expressed at the barrier sites, whereas CD147, cyclophilins, CD26 and their interaction partners are present in immune cells and in epithelium. Expression of **A)** *ACE2*, **B)** *TMPRSS2*, **C)** *BSG*, **D)** *PPIA*, **E)** *PPIB*, **F)** *S100A9*, **G)** *SLC16A7*, **H)** *SLC2A1*, **I)** *CD44*, **J)** *ITGB1*, **K)** *NFATC3*, **L)** *DPP4* genes in *in vitro* Air Liquid Interface (ALI) – differentiated human primary bronchial epithelial cells (n=10) and in *ex vivo* human primary bronchial biopsies (n=5), bronchoalveolar fluid cells (n=5), whole blood (n=5), neutrophils (n=4), classical monocytes (n=4), plasmocytoid dendritic cells (n=4), group 1, 2 and 3 innate lymphoid cells (n=6 per group), natural-killer cells (n=4), naïve CD4^+^ T cells (n=4), terminal effector CD4^+^ T cells (n=2), naïve CD8^+^ T cells (n=4), effector memory CD8^+^ T cells (n=4), naïve B cells (n=4), plasmablasts (n=4) and skin biopsies (n=6) from healthy adults. Data obtained from *in vitro* approaches are highlighted in red. Names of the proteins encoded by analysed genes are stated in the brackets. HBECS, human bronchial epithelial cells; Bronch. biop., bronchial biopsy; BAL, bronchoalveolar fluid cells; Class. monocytes, classical monocytes; pDCs, plasmocytoid dendritic cells; ILC1, group 1 innate lymphoid cells; ILC2, group 2 innate lymphoid cells, ILC3, group 3 innate lymphoid cells; NK, natural killer cells; naïve CD4^+^, naïve CD4^+^ T cells; term. Eff. CD4^+^, terminal effector CD4^+^ T cells; naïve CD8^+^, naïve CD8^+^ T cells, eff. mem. CD8^+^, effector memory CD8^+^ T cells.

In contrast to *ACE2*, we observed a prominent expression of CD147 (*BSG*) in epithelial and innate and adaptive immune cells in the lung (BAL), in the skin and in circulation (whole blood, neutrophils, classical monocytes, pDCs, *ex vivo* sorted ILC, NK, naïve CD4^+^ T cells, terminal effector CD4^+^ T cells, naïve CD8^+^ T cells, effector memory CD8^+^ T cells, naïve B cells and plasmablasts) (Figure 1C, Figures S1-S3). Interestingly, *ACE2* and CD147 (*BSG*) are not co-expressed in bronchial biopsy (Figure S2). High expression of CD147 (*BSG*) in primary human bronchial epithelial cells suggests that these cells having more than one receptor for SARS-CoV-2 entry could be infected in various conditions. Granulocytes and macrophages are the main source of CD147 (*BSG*) expression^49^, which may explain high CD147 (*BSG*) expression in the whole blood and BAL observed in our data. Presence of CD147 (*BSG*) on innate and adaptive memory immune cells may indicate the importance of this receptor in systemic inflammatory response and development of immunological memory.

Since it is not clear yet how exactly the process of infection via CD147 (*BSG*) takes place in case of SARS-CoV-2, we examined the presence and co-expression of extracellular partners of CD147 (*BSG*), which are implicated in the infection with other viruses using CD147 (*BSG*) for cell entry, including SARS-CoV, HIV-1, measles and others^11–13^. All investigated cell populations showed high expression of cyclophilin A and B (*PPIA* and *PPIB*) (Figure 1D and E, Figure S1). Interestingly, cyclophilin A was highly expressed in all ILC. In addition, cyclophilin B was highly expressed in pDCS, ILC and plasmablasts. *S100A9* was highly expressed not only in bronchial epithelium, but also in the whole blood, neutrophils, classical monocytes and, surprisingly, in naïve B cells (Figure 1F and Figure S1).

Next, we examined transmembrane and intracellular partners of CD147 (*BSG*), because its cytoplasmic tail, bound to many transmembrane partners, is essential in the entry of other viruses^29^. We observed that monocarboxylate transporters, such as *SLC16A7* (MCT2) and *SLC2A1* (GLUT1) are highly expressed not only at the barrier sites, but also in ILC (MCT2), pDCs, NK cells, T cells and B cells (MCT2, GLUT1) (Figure 1G and H). *CD44* is highly expressed in majority of cells, with the exceptionally high expression in BAL, classical monocytes, naïve CD4^+^ T cells, terminal effector CD4+ T cells, naïve CD8^+^ T cells and effector memory CD8^+^ T cells (Figure 1I). *ITGB1* (Integrin β-1) and other integrins are highly expressed in airway epithelial cells, but also other cells expressed them (Figure 1J and Figure S1). NF-ATc1-3 (*NFATC1, NFATC2, NFATC3*) were expressed predominantly in CD4^+^ and CD8^+^ T and naïve B cells, whereas expression of *NFAT4* and *NFAT5* was higher in the lung and skin (Figure 1K and Figure S1).

Finally, we looked at the expression of CD26 (*DPP4*). Notably, it was expressed highly in all ILC, and in naïve CD4^+^ and CD8^+^ T cells, as well as in pDCs and effector memory CD8^+^ T cells (Figure 1L and Figure S1).

### Distinct expression profiles of ACE2-, CD147-, and CD26-related genes in PBMCs and T cells of children, adolescents and adults

Clinical evidence from COVID-19 patients worldwide clearly demonstrate an association between disease severity and morbidity with age^1^. In China, less than 1% of the SARS-CoV-2 positive cases were in children younger than 10 years of age^50,51^. Therefore, we analyzed expression patterns of ACE2-, CD147-, and CD26-related genes in peripheral blood mononuclear cells (PBMCs) from infants, young children, adolescents and adults. While these comparisons can be limited due to the different distribution of immune subsets in the blood of children and adults^52^ and different ethnicity of children (African American from South Africa and Tanzania) compared to adolescents and adults (Caucasians), we observed that there are different patterns of expression for the majority of analyzed genes (Figure 2A). We observed that CD147 (*BSG*) was highly expressed in PBMCs of children, whereas its expression was lower in adolescents and in adults. Similarly, to CD147 (*BSG*), cyclophilins B (*PPIB*), *S100A9, SLC3A2* (CD98), *SLC16A1* (MCT1), *NFATC4* were less expressed in adults. On the other hand, *SLC16A3* (MCT4), *SLC16A7* (MCT2), *NFATC2, NFAT5, NFATC1, NFATC3*, integrins (*ITGB1*) were expressed on higher level in adults. Older children and adolescents showed intermediary gene expression between children and adults.

**Figure 2.**
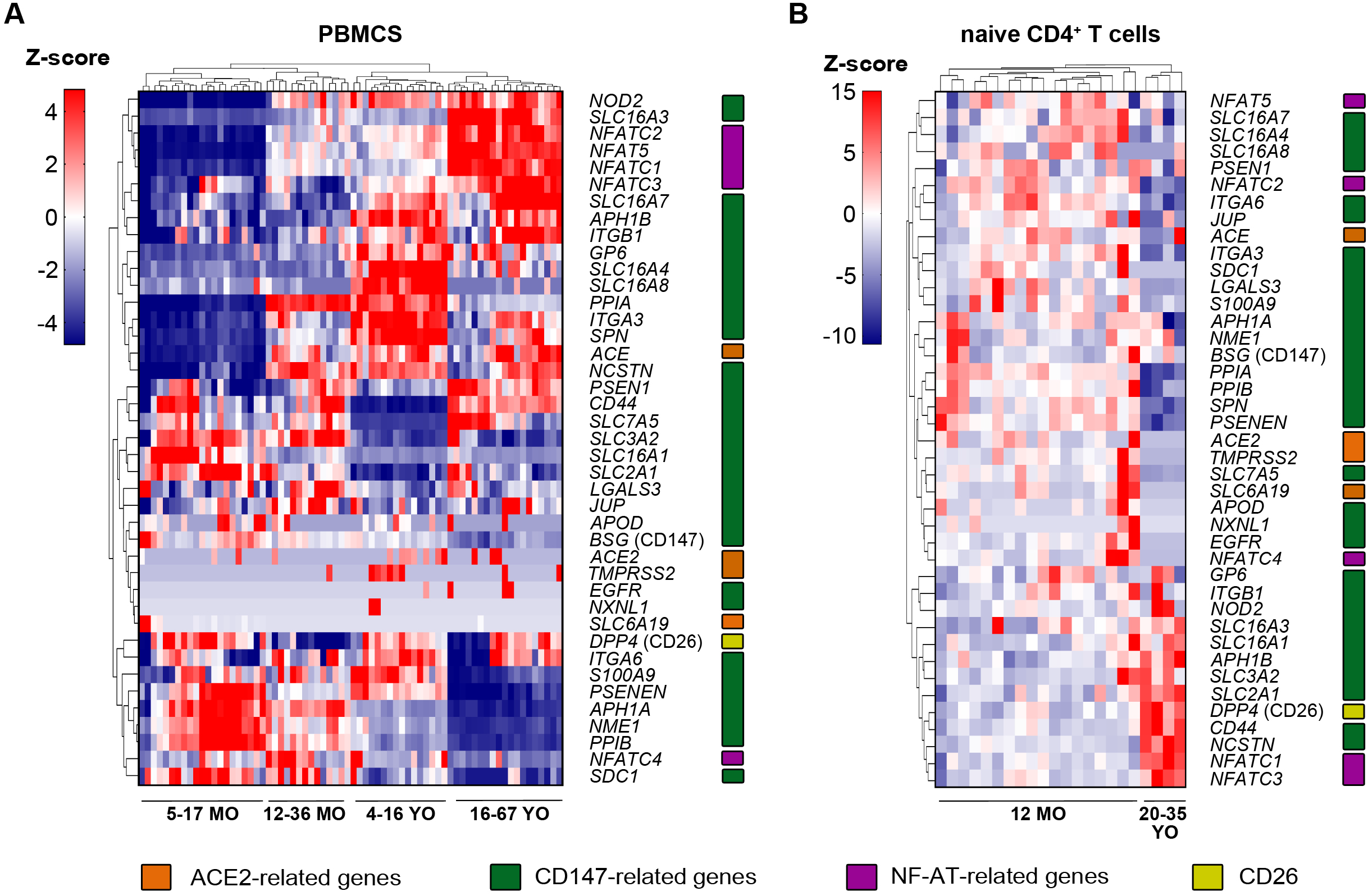
Distinct expression profile of ACE2-, CD147-, and CD26-related genes in PBMCs and T cells of children and adults. **A)** Expression of ACE2-, CD147-, NFAT- and CD-26-related genes in the primary human PBMCs in healthy children aged 5-17 months (n=21), 12-36 month (n=14), 4-16 years (n=16) and healthy adults aged 16-67 years (n=19). **B)** Expression of ACE2-, CD147-, NFAT- and CD-26-related genes in the primary human naïve CD4^+^ T cells from 12 months old healthy children (n=18) and 20-35 years old healthy adults (n=4). All heatmaps display normalized gene expression across the groups (rows normalization). Color-coding represents gene families related with ACE2 (orange), CD147 (green), NF-ATs (purple) and CD26 (yellow). MO, months old, YO, years old, PBMCs, peripheral blood mononuclear cells.

Next, we compared ACE2-, CD147- and CD26-related gene expression profile in the purified naïve CD4^+^ T cells between young children and adults (Figure 2B). Low expression of ACE2-related genes was observed in both children and adults. Several CD147-related genes showed higher expression in adults, including *CD44*, certain MCTs, *SLC3A2* (CD98), *SLC2A1* (GLUT1), *NFATC1* and *NFATC3*. Also, CD26 (*DPP4*) was expressed at a higher level in adults. In contrast, other CD147-related genes were expressed at lower levels in adults, such as cyclophilins A and B (*PPIA* and *PPIB), S100A9, SLC7A5* (CD98), *LGALS3*, integrins (*ITGA6, ITGA3*) and *NFATC2*.

### Asthma, COPD, hypertension, smoking, obesity and gender show different expression profiles of ACE2-, CD147-, and CD26-related genes

We analyzed expression of ACE2-, CD147-and CD26-related genes from different cells and tissues in various comorbidities and risk factors, which have been shown, or are suspected to predispose to SARS-CoV-2 infection and/or COVID-19 progression. Upper and lower airways are the initial entry of SARS-CoV-2, thus we first analyzed gene expression in the HBECs and the bronchial biopsies from patients with asthma or COPD as compared to the non-diseased controls. In our cohorts, we did not see any significant difference in *ACE2* expression in HBECs or bronchial biopsy between control, asthma and COPD patients (Figure 3A and B). However, in bronchial biopsies *ACE2* expression was higher in smokers (Figure 3B). In HBECs, we observed higher expression of *TMPRSS2* in asthma (Figure 3A), raising the possibility that in asthmatic airways the cleavage of spike protein of SARS-CoV-2 might be more efficient. We also observed a trend of increased expression of CD147 (*BSG*) in HBECs (Figure 3A) and in bronchial biopsies (Figure 3B) from COPD patients. Additionally, glucose transporter GLUT1 (*SLC2A1*), integrin α-3 (*ITGA3*) and galectin-3 (*LGALS3*) were higher expressed in HBECs, whereas *SLC7A5* (CD98), integrins α-3 (*ITGA3*) and α-6 (*ITGA6*) were higher expressed in the bronchial biopsies of COPD patients. *CD44* and *APOD* were higher expressed in asthma (Figure 3A, B and Figures S4, S5). In bronchial biopsies, ACE2-, CD147- and CD26-related genes showed similar cluster of co-expression in control and asthma (Figure S2 and S6). Interestingly, *ACE2* co-expressed with *PPIB* and *NME1* in asthma, which was not found in controls (Figure S6). Taken together, airway epithelium in asthma and COPD showed a gene signature that potentially can facilitate SARS-CoV-2 entry and enhance internalization after receptor binding.

**Figure 3.**
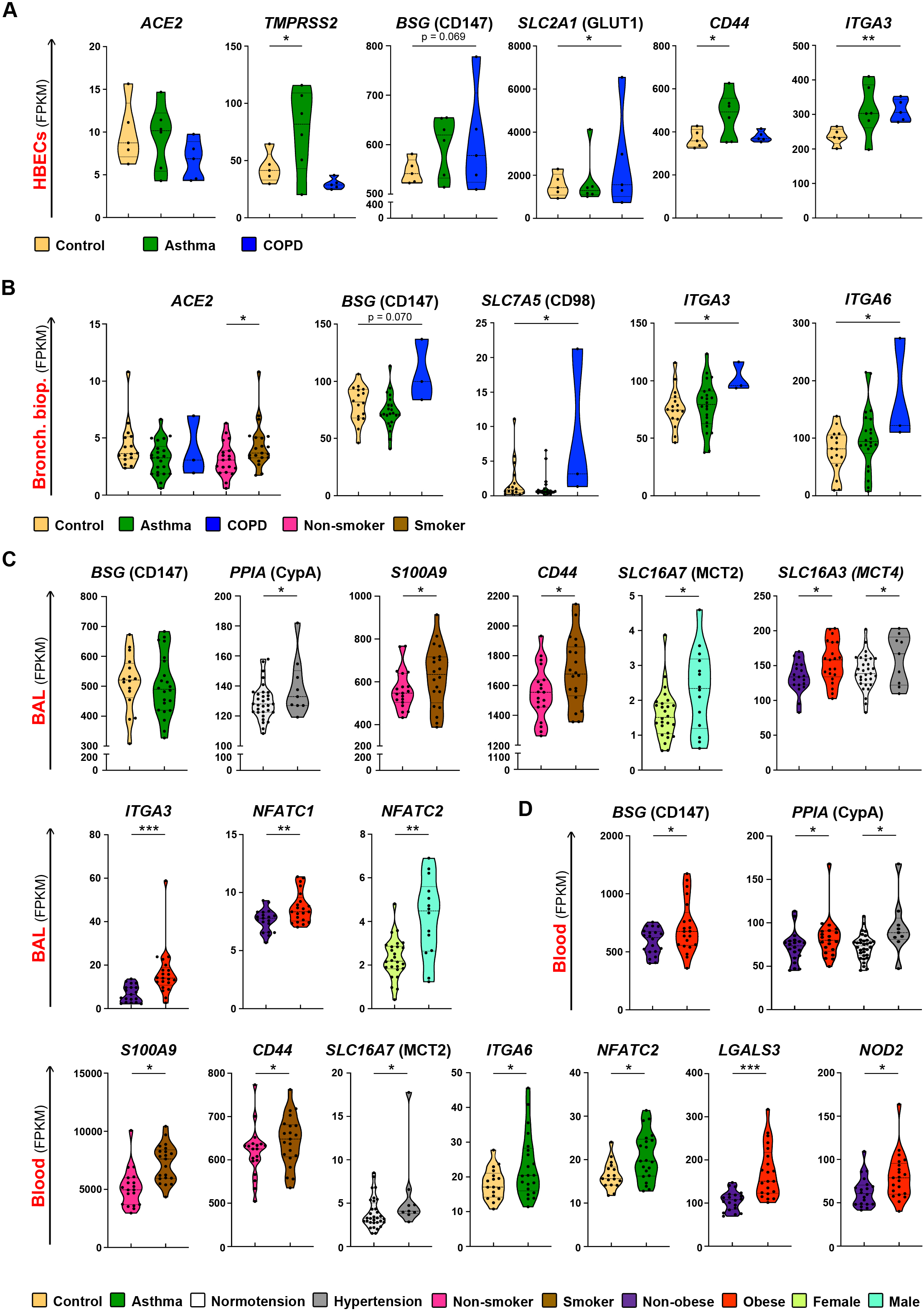
Asthma, COPD, hypertension, smoking, obesity and gender is associated with differential expression of ACE2-, CD147-, and CD26-related genes in immune cells and tissues. **A)** Differential expression of *ACE2, TMPRSS2, BSG, SLC2A1, CD44* and *ITGA3* genes in *in vitro* Air Liquid Interface (ALI) – differentiated human primary bronchial epithelial cells from non-diseased controls (n=5), asthma (n=6) and COPD (n=5) patients. **B)** Differential expression of *ACE2, BSG, SLC7A5, ITGA3, ITGA6* genes in bronchial biopsies from non-diseased controls (n=16), patients with asthma (n=22) and COPD (n=3), or in comparison of smokers (n=21) with non-smoking individuals (n=19). **C)** Differential expression of *BSG*, *PPIA*, *S100A9*, *CD44*, *SLC16A7*, *SLC16A3*, *ITGA3*, *NFATC1*, *NFATC2* genes in the bronchoalveolar fluid (BAL) from the control individuals (n=16), patients with asthma (n=22) and COPD (n=2), or in comparison of hypertensive (n=9) with normotensive (n=31) individuals; smokers (n=20) with non-smokers (n=19); obese (n=21) with non-obese (n=19); and males (n=26) with females (n=14). **D)** Differential expression of *BSG*, *PPIA, S100A9, CD44, SLC16A7, ITGA6, NFATC2, LGALS3* and *NOD2* genes in the whole blood of non-diseased controls (n=17), patients with asthma (n=21) and COPD (n=3), or in comparison of hypertensive (n=9) with normotensive (n=32) individuals; smokers (n=21), with non-smokers (n=19); obese (n=21) with non-obese individuals (n=20); and males (n=27) with females (n=14). Names of the proteins encoded by analyzed genes are stated in the brackets. *p < 0.05, **p < 0.01, ***p < 0.001, ****p< 0.0001. HBECS, human bronchial epithelial cells; Bronch. biop., bronchial biopsy; BAL, bronchoalveolar fluid cells; COPD, chronic obstructive pulmonary disease.

Next, we analyzed BAL and whole blood gene expression in patients with asthma and controls (Figure 3 C, D and Figure S7, S8). Unfortunately, we have access to the limited number of BAL samples from COPD patients. In BAL, which reflects local lung immune microenvironment, and in blood, reflecting systemic immune responses, CD147 (*BSG*) was expressed equally high in asthma patients and in controls (Figure 3C, Figure S7). In blood, patients with asthma had also higher expression of integrin *ITGA6* and *NFATC2*, as compared to controls (Figure 3D). There was a greater abundance in the cluster of co-expressed genes in asthma in BAL and especially in blood (Figure S9, S10). In asthma patients, *BSG* (CD147) showed co-expression with *SLC7A5* (CD98) in blood, *DPP4* (CD26) also co-expressed with *CD44* and *ITGA6* (Figure S10).

Next, we analyzed the ACE2-, CD147- and CD26-related gene expression in our controls, asthma and COPD patients according to the additional clinical features, which have been reported as a risk factor comorbidity for COVID-19, such as hypertension, smoking, gender and obesity^23,53^. We did not see any major differences in the gene expression in the bronchial biopsies based on these features, except higher *ACE2* expression in smokers (Figure 3B, Figure S5). However, we noted several important differences in the BAL and in the whole blood (Figure 3 C, D and Figure S7, S8). Subjects with hypertension had increased expression of cyclophilin A in both the BAL and blood, as well as upregulated expression of MCT4 (*SLC16A3*), *APH1A* and *PSENEN* in the BAL and MCT2 (*SLC16A7*) in the blood (Figure 3 C, D and Figure S7, S8). In case of smoking, expression of *S100A9* and *CD44* was elevated in both BAL (Figure 3C, Figure S7) and blood (Figure 3D, Figure S8). Obesity was an important factor leading to the significant changes in the BAL and blood. We observed that MCT4 (*SLC16A3*), integrin *ITGA3, NFATC1* and *PSEN1* were more expressed in the BAL of obese individuals (Figure 3C, Figure S7), whereas CD147 (*BSG*), *PPIA, LGALS3* and *NOD2* were more expressed in their blood (Figure 3D). Finally, regarding gender in our cohort, MCT2 (*SLC16A7*), CD98 (*SLC7A5*), *NFATC2* were higher expressed in the BAL of male subjects (Figure 3C, Figure S7).

### Expression of CD147-related genes correlates with BMI and age in the BAL and blood

Since obesity and age were important variables, we additionally correlated these two variables with the expression of ACE2-, CD147- and CD26-related genes in the bronchial biopsy, BAL and blood of our adult non-diseased and diseased cohorts. We did not find any significant correlations between BMI and age and the receptor-related gene expression in the bronchial biopsies. However, we noted that the expression of *SLC16A3* (MCT4), *ITGA3, LGALS3* in BAL positively correlated with BMI (Figure 4A-C) and the expression of *CD44* positively correlated with the age of the subjects (Figure 4D). Interestingly, whole blood expression of CD147 (*BSG*), *PPIA, S100A9, CD44* and *LGALS3* correlated positively with the BMI, whereas *SLC16A3* positively correlated with age (Figure 4 E-J). In summary, these findings suggest that higher BMI and older age lead to higher expression of CD147-related genes on immune cells, but not on the barrier cells, which potentially can influence the development and the course of COVID-19.

**Figure 4.**
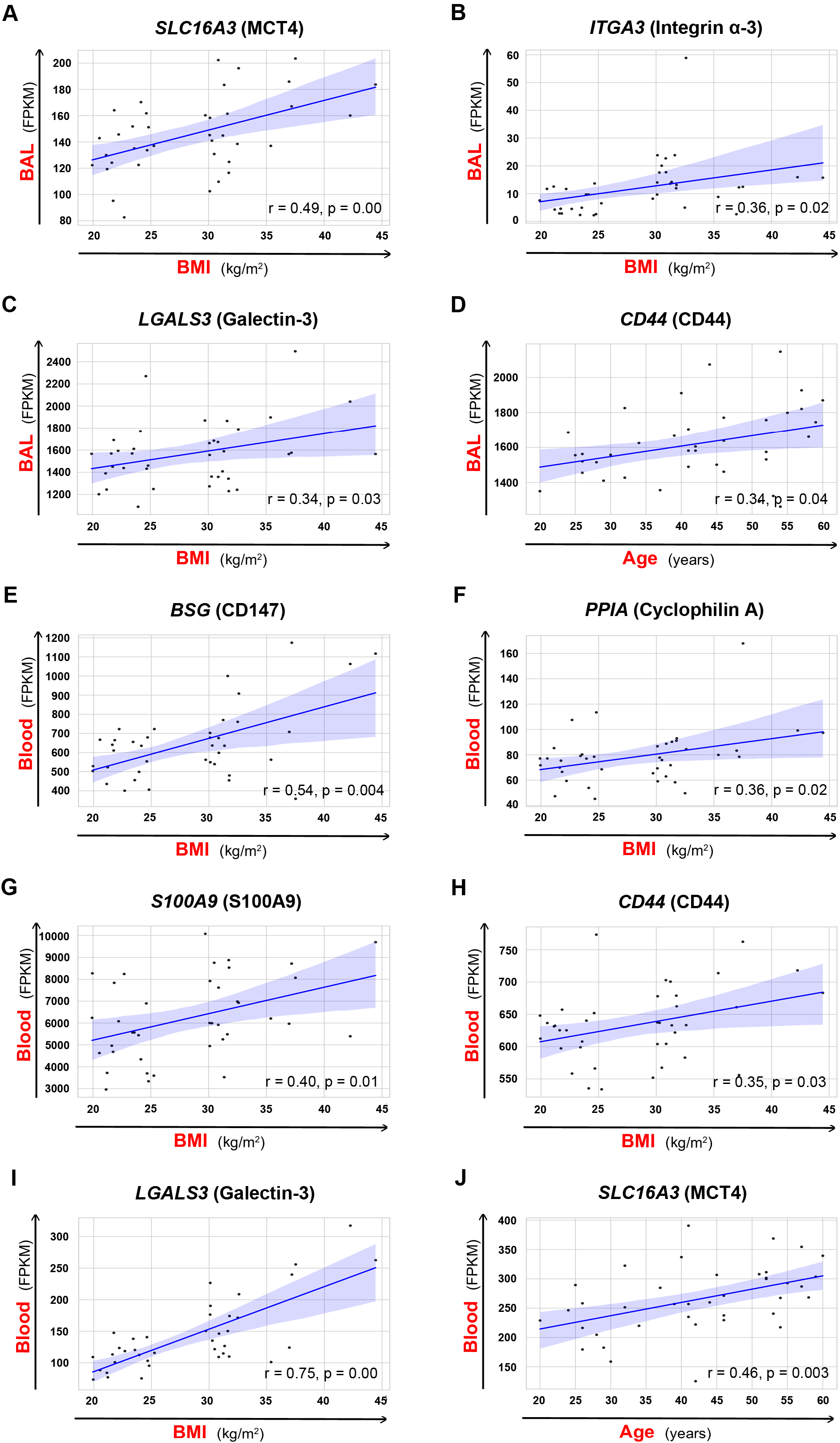
Expression of certain CD147-related genes correlates with BMI and age in the BAL and blood. Correlation of **A)** *SLC16A3* expression and BMI, **B)** *ITGA3* expression and BMI, **C)** LGALS3 expression and BMI, and **D)** *CD44* expression and age in the bronchoalveolar fluid (BAL). Correlation of **E)** *BSG* expression and BMI, **F)** *PPIA* expression and BMI, **G)** *S100A9* expression and BMI, **H)** *CD44* expression and BMI, **I)** *LGALS3* expression and BMI, **J)** *SLC16A3* and age in the whole blood. Spearman correlation coefficient (r) was calculated, with the threshold of significance set to p = 0.05. Names of the proteins encoded by analyzed genes are stated in the brackets. BAL, bronchoalveolar fluid cells; BMI, body-mass index.

### Eczema lesional skin in patients with atopic dermatitis present unique ACE2- and CD147-related gene expression profile

As we found a high expression of ACE2-, CD147- and CD26-related genes in the skin biopsies of healthy subjects, and since COVID-19-related skin lesions have been recently reported^54^, we also analyzed the biopsies of healthy and atopic dermatitis (AD) skin. Interestingly, in the lesional skin as compared to non-lesional skin of the same subjects, many CD147-related genes showed higher expression, including cyclophilins (*PPIA* and *PPIB), S100A9, CD44*, MCTs, *SLC7A5* (CD98) and integrins (Figure 5).

**Figure 5.**
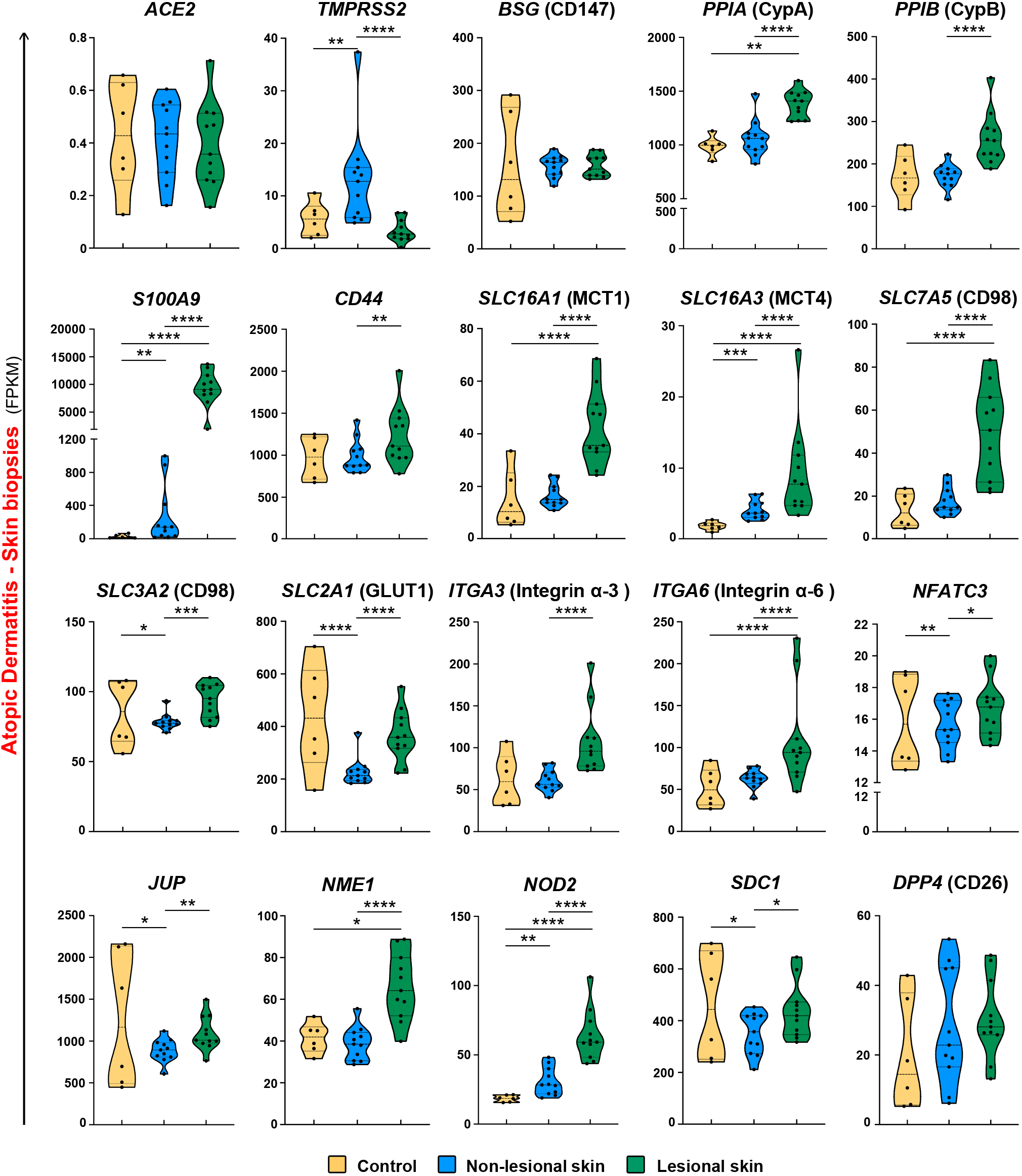
Unique expression profile of ACE2- and CD147-related genes in lesional skin in patients with atopic dermatitis. **A)** Expression of *ACE2, TMPRSS2, BSG, PPIA, PPIB, S100A9, CD44, SLC16A1, SLC16A3, SLC7A5, SLC3A2, SLC2A1, ITGA3, ITGA6, NFATC3, JUP, NME1, NOD2, SDC1 and DPP4* genes in the skin of healthy controls (n=6) and in the lesional (n=11), and non-lesional (n=11) skin of atopic dermatitis patients. Names of the proteins encoded by analyzed genes are stated in the margins. *P < 0.05, **P < 0.01, ***P < 0.001, ****P < 0.0001.

## Discussion

COVID-19 pandemic is developing at such a pace that extraordinary actions are being initiated to learn quickly about its biology, transmission and potential means of prevention and treatment. In this spirit, we first performed an extensive literature and database search and curated a list of proven and potential receptors for SARS-CoV-2 and interaction partners, whose expression in different tissues and cells might be involved in the course of immunological response in COVID-19 (Figure 6)^55^. Next, we analyzed these genes in the broad spectrum of cell types and tissues in healthy controls to evaluate the level of their expression and their co-expression profiles, as well as evaluated their expression in healthy children, adolescents and adults. While interesting associations were observed to be age-dependent, these findings must be further explored and repeated due to the possibility of batch-specific systematic variations in the gene expression values between in-house and public datasets. Despite our efforts in applying identical data analysis workflow for the datasets produced by our group and those available in public repositories, different sequencing facilities and experiment protocols can lead to altered gene expression values, especially for genes with extreme gene length and G/C content. Finally, we analyzed gene expression in adults with known and potential COVID-19 comorbidities and risk factors such as COPD, asthma, hypertension, obesity, smoking, male gender and AD. These conditions can be directly compared, as they were performed internally in the context of same project.

**Figure 6.**
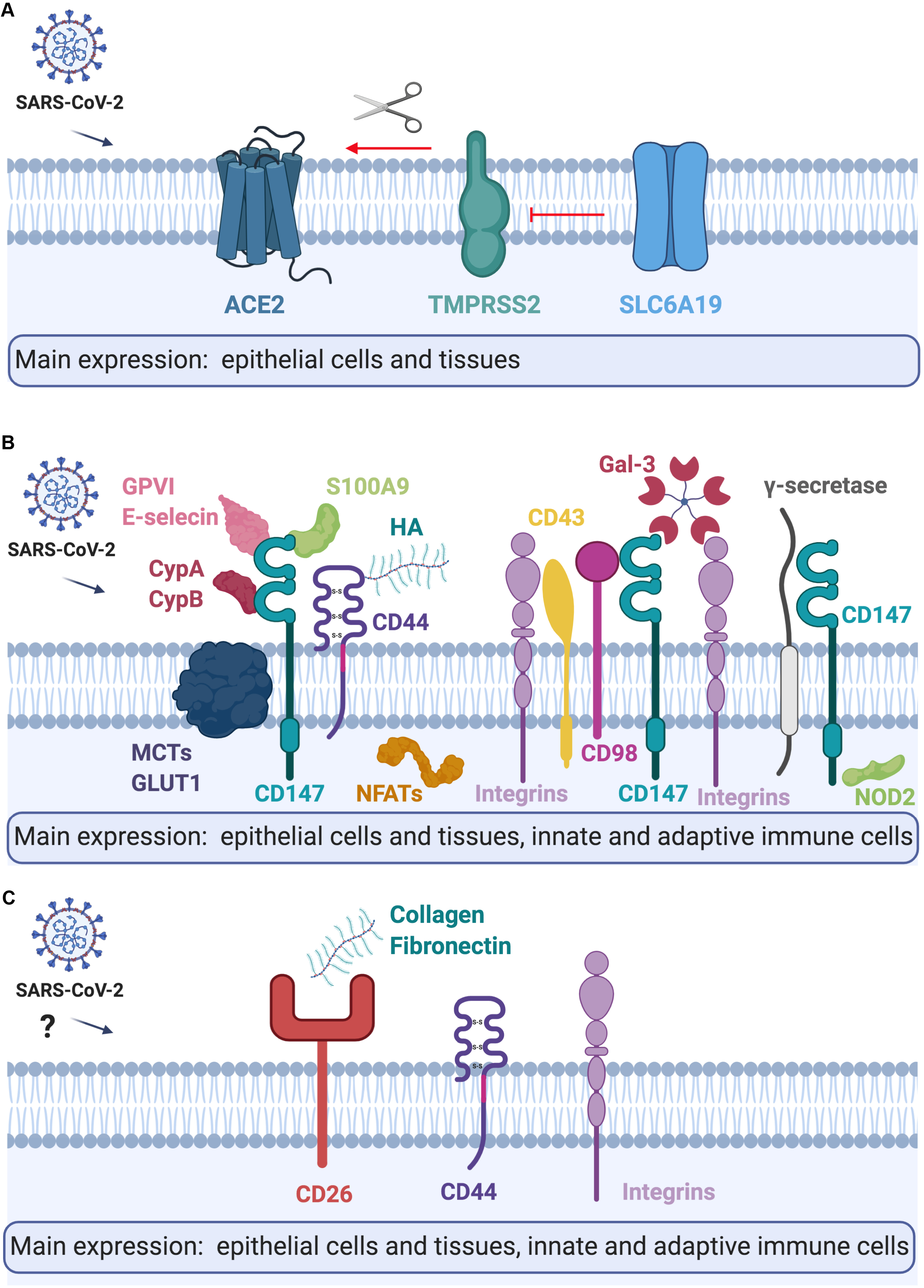
Summary of the tissue and cellular expression, and models of **A)** ACE2, **B)** CD147, **C)** CD26 and their interaction partners. Please refer to the text for further details. CypA, Cyclophilin A; CypB, Cyclophilin B; HA, Hyaluronic acid; Gal-3, Galectin 3; MCTs, Monocarboxylate transporters. Figure created with BioRender.com.

ACE2 is a receptor for SARS-CoV-2^3^, whereas TMPRSS2 is a transmembrane host protease, which cleaves the viral spike protein thus facilitating virus fusion to the cellular membranes process (S)^3,56^. SLC6A19, physiologically a neutral amino acid transporter, potentially can block the access of TMPRSS2 to ACE2 and subsequently reduce active infection^4,6^. Our results, in agreement with recent reports^6,57,58^, indicate that airway epithelium has high ACE2 and TMPRSS2 co-expression, and no expression of protective SLC6A19 and thus might be highly susceptible for SARS-CoV-2 infection. In addition, it has been shown recently that ACE2 is highly expressed in naso- and oropharynx, which are the sites of active SARS-CoV-2 replication and a main source of infectious particles^6,57,58^. Interestingly, we observed high expression of ACE2 and TMPRSS2 in bronchial airway epithelium: in HBECs ALI cultures *in vitro*, as well as *in vivo* in humans in bronchial biopsies, indicating that they can be as important in initiation and progression of COVID-19, as recently shown for type II pneumocytes^58^. We have not observed the expression of SLC6A19 in any of the analyzed cells and tissues, yet it is known to be expressed is intestine^59^. Interestingly, in contrast to Leung and colleagues^60^, we have not seen higher expression of ACE2 in HBECs or bronchial biopsy of patients with COPD, but our sample size was very limited, and potentially a bronchial biopsy site was different, which might be the reason of this inconsistency. However, consistently with this report^61^, we observed higher expression of ACE2 in the bronchial biopsy of current smokers, which might indicate that current smoking status, might be a stronger factor of increased ACE2 expression in airway epithelium than COPD per se. On the other hand, ACE2 and the angiotensin II type 2 receptor (AT2) were reported to protect from severe acute lung injury in mice^62^, therefore an importance of ACE2 in SARS-CoV-2 infection and COVID-19 progression should be further explored. In addition, in HBECs of patients with asthma, we observed higher expression of TMPRSS2, which increases the possibility of SARS-CoV-2 cleavage in asthmatic bronchi. Even though initially asthma was not reported to be a significant comorbidity for COVID-19, more observations from Europe and the US seem to show otherwise^63^. It might be related to the heterogeneity of asthma endotypes type 2 asthma (allergic) and non-type 2 (non-allergic). Allergen exposure, allergic sensitization and high IgE lead to lower ACE2 expression in the nasal and bronchial epithelium of asthma patients^64^. Thus, it is possible that ACE2 expression in the airways of allergic patients, even if slightly protective in terms of infection with SARS-CoV-2, might also predispose to faster progression of COVID-19 to more severe forms, especially in case of higher TMPRSS2 expression.

We found that immune cells do not express ACE2, TMPRSS2 or SLC6A19, which has also been observed by others^2,5^ Importantly though, we found that CD147 and its extracellular agonists and transmembrane partners are highly expressed in innate and adaptive immune cells in the lungs and in the periphery, suggesting that they should be further investigated in SARS-CoV-2 spread and COVID-19 pathology^9^. Here, we report high expression of CD147 in both, epithelial tissues and innate and adaptive immune cells. Potentially, epithelial cells, macrophages, monocytes, ILCs, NK cells, T cells and B cells can be infected in the lungs or can carry SARS-CoV-2 from infected epithelial cells via CD147 and participate in the local and systemic spread of the virus, and in exaggerated immune response^65^.

We found here that CD147 is slightly higher expressed in the HBECs and bronchial biopsy of COPD patients, as well as it is higher expressed in the blood of obese individuals. Obesity is one of the main comorbidities reported in patients with severe COVID-19^63^, and since CD147 expression in the whole blood correlates positively with BMI in our cohort, it certainly needs further attention in relation to COVID-19. Importantly, CD147 expression is upregulated by high glucose concentrations^66^, which might reflect its correlation with obesity, and potentially also with diabetes, another very important COVID-19 comorbidity. Additionally, extracellular ligands and transmembrane partners of CD147 participate in the progress of other viral infections^12^ in variety of mechanisms. Viruses integrate cyclophilin A and cyclophilin B in their virions and further can bind and either infect cells via 147 or activate CD147 signaling^11–13^. We found that expression of both cyclophilins (*PPIA* and *PPIB*) is increased in subjects with hypertension and obesity and *PPIA* correlated positively with BMI in the whole blood, which is confirmed in other studies^67^. In addition, nsp1 protein of SARS-CoV binds to CD147 via cyclophilins and reduces interferon responses^25^ in infected cells. It also inhibits NF-ATs translocation activation in T cells, leading to suppression of immune responses^25,68^. Both mechanisms might be more potent in obese patients, especially with hypertension, due to higher expression of both cyclophilins on the periphery. Also, the other extracellular ligand of CD147-S100A9 correlated positively with BMI. CD147 transmembrane partners such as *LGALS3*, MCT4 (*SLC16A3*), MCT2 (*SLC16A7*), *ITGA3* and *NFATC1* were elevated in the blood or BAL of obese or hypertensive individuals, as well as the expression of *LGALS3, S100A9* and *CD44* correlated positively with BMI. Importantly in individuals who are current smokers *S100A9* and *CD44* are elevated in the BAL and in the blood. MCTs regulate the transport of lactate, pyruvate and ketone bodies across the cell membrane^69^. Galectin-3 (*LGALS3*) via CD147 signaling is responsible for disrupting cell-cell contact at the epithelial barriers^70^. CD44 in acute inflammation triggered by hyaluronic acid lead to the cell activation and release of many proinflammatory cytokines whereas in the longer terms induce lung fibrosis^71,72^. Interestingly, it has been shown that patients who recovered from SARS-CoV have spike protein-specific memory cytotoxic T cells, which express CD44^73^. Thus, our results suggest a possible role of CD147 and its extracellular and intracellular ligands in COVID-19 pathogenesis via regulation of cell metabolism, motility and activation, especially in patients with comorbidities. A clinical trial with anti-CD147 in COVID-19 patients seems to confirm its pleiotropic role in SARS-CoV-2 infection, but it requires further research to investigate the involved mechanisms and molecules.

Finally, expression of ACE2/TMPRSS2, CD147- and CD26-related genes in the skin biopsies highlights the potential roles of these molecules in various viral diseases. Recent studies reported that many COVID-19 cases developed nonpruritic, erythematous rashes, urticaria or varicella-like lesions^54,74^, but so far it is not known if they are a place of viral replication or just a local reaction to systemic infection. Similarly, not much is yet known about the potential connections between AD and the course of COVID-19^75^. Yet, severe and untreated AD is a known risk factor for disseminated viral skin disease^76^. Therefore, our observations indicating higher expression of cyclophilins, CD44, S100A9, integrins, MCTs and other CD147-related genes require further studies about their role in the course of SARS-CoV-2 infection in healthy subjects and in AD patients.

CD26 (*DPP4*) is another receptor important in coronavirus infection, described in MERS-CoV, potentially recognizing SARS-CoV-2^21,77^. Similar to CD147, CD26 was expressed in all investigated cell types, except B cells. ILCs showed highest expression of DPP4 from all analyzed cells, which is in agreement with recent findings^78^. However, not much is known about the variable expression of CD26 among ILC1, ILC2 and ILC3. Potentially, a high level of CD26 on the surface of ILCs in the lung may be another mechanism of systemic spread of the virus.

The differential profile of ACE2-, CD147- and CD26-related gene expression in healthy subjects of different ages need to be interpreted with caution due to the distinct origins of these populations, as well as potential experimental bias (even if we performed normalization strategies, similar bioinformatic pipelines and cautious assessment of raw data quality, prior to analysis). Nevertheless, our observations are largely in agreement with other recent findings correlating gene expression in the whole blood and PBMCs with age^79^. Having this in mind, it is intriguing to observe that expression of NOD2, some MCTs, integrins and NFATs seemed to increase with age, whereas the expression of genes coding γ-secretase complex is decreasing with age in PBMCs. CD44 and MCT4 also correlated with age in BAL and blood, respectively. Moreover, some of the differences between children and adults, such as higher expression of CD26, CD44, GLUT1 (*SLC2A1*) and specific NFATs in adults were also observed in naïve purified CD4^+^ T cells. This requires further investigation, but differences in expression pattern may be related to the striking differences in the morbidity of SARS-CoV-2 between children and adults^80^.

## References

1. Zhang JJ, Dong X, Cao YY, et al. Clinical characteristics of 140 patients infected with SARS-CoV-2 in Wuhan, China. Allergy. 2020.

2. Wu A, Peng Y, Huang B, et al. Genome Composition and Divergence of the Novel Coronavirus (2019-nCoV) Originating in China. Cell Host Microbe. 2020;27(3):325–328.

3. Hoffmann M, Kleine-Weber H, Schroeder S, et al. SARS-CoV-2 Cell Entry Depends on ACE2 and TMPRSS2 and Is Blocked by a Clinically Proven Protease Inhibitor. Cell. 2020;181(2):271–280 e278.

4. Yan R, Zhang Y, Li Y, Xia L, Zhou Q. Structure of dimeric full-length human ACE2 in complex with B<sup>0</sup>AT1. bioRxivpreprint. 2020:2020.2002.2017.951848.

5. Ziegler C, Allon S, Nyquist S, et al. SARS-CoV-2 Receptor ACE2 is an Interferon-Stimulated Gene in Human Airway Epithelial Cells and Is Enriched in Specific Cell Subsets Across Tissues. SSRN Electronic Journal. 2020.

6. Wu C, Zheng, M. Single-cell RNA expression profiling shows that ACE2, the putative receptor for COVID-2019, has significant expression in nasal and mounth tissue and is co-expressed with TMPRSS2 and not co-expressed with SLC6A19 in the tissues. PREPRINT (Version 1) available at Research Square. 12 March 2020.

7. Chu H, Zhou J, Wong BH, et al. Middle East Respiratory Syndrome Coronavirus Efficiently Infects Human Primary T Lymphocytes and Activates the Extrinsic and Intrinsic Apoptosis Pathways. J Infect Dis. 2016;213(6):904–914.

8. Gu J, Gong E, Zhang B, et al. Multiple organ infection and the pathogenesis of SARS. J Exp Med. 2005;202(3):415–424.

9. Wang K, Chen W, Zhou Y-S, et al. SARS-CoV-2 invades host cells via a novel route: CD147-spike protein. bioRxiv. 2020:2020.2003.2014.988345.

10. Wang X, Xu W, Hu G, et al. SARS-CoV-2 infects T lymphocytes through its spike protein-mediated membrane fusion. Cell Mol Immunol. 2020.

11. Chen Z, Mi L, Xu J, et al. Function of HAb18G/CD147 in invasion of host cells by severe acute respiratory syndrome coronavirus. J Infect Dis. 2005;191(5):755–760.

12. Pushkarsky T, Zybarth G, Dubrovsky L, et al. CD147 facilitates HIV-1 infection by interacting with virus-associated cyclophilin A. Proc Natl Acad Sci U S A. 2001;98(11):6360–6365.

13. Watanabe A, Yoneda M, Ikeda F, Terao-Muto Y, Sato H, Kai C. CD147/EMMPRIN acts as a functional entry receptor for measles virus on epithelial cells. J Virol. 2010;84(9):4183–4193.

14. Zhang MY, Zhang Y, Wu XD, et al. Disrupting CD147-RAP2 interaction abrogates erythrocyte invasion by Plasmodium falciparum. Blood. 2018;131(10):1111–1121.

15. Bian H, Zheng Z-H, Wei D, et al. Meplazumab treats COVID-19 pneumonia: an open-labelled, concurrent controlled add-on clinical trial. medRxiv. 2020:2020.2003.2021.20040691.

16. Muramatsu T. Basigin (CD147), a multifunctional transmembrane glycoprotein with various binding partners. J Biochem. 2016;159(5):481–490.

17. Yurchenko V, Constant S, Eisenmesser E, Bukrinsky M. Cyclophilin-CD147 interactions: a new target for anti-inflammatory therapeutics. Clin Exp Immunol. 2010;160(3):305–317.

18. Hibino T, Sakaguchi M, Miyamoto S, et al. S100A9 is a novel ligand of EMMPRIN that promotes melanoma metastasis. Cancer Res. 2013;73(1):172–183.

19. Kato N, Yuzawa Y, Kosugi T, et al. The E-selectin ligand basigin/CD147 is responsible for neutrophil recruitment in renal ischemia/reperfusion. J Am Soc Nephrol. 2009;20(7):1565–1576.

20. Seizer P, Borst O, Langer HF, et al. EMMPRIN (CD147) is a novel receptor for platelet GPVI and mediates platelet rolling via GPVI-EMMPRIN interaction. Thromb Haemost. 2009;101(4):682–686.

21. Huang W, Luo WJ, Zhu P, et al. Modulation of CD147-induced matrix metalloproteinase activity: role of CD147 N-glycosylation. Biochem J. 2013;449(2):437–448.

22. Vankadari N, Wilce JA. Emerging WuHan (COVID-19) coronavirus: glycan shield and structure prediction of spike glycoprotein and its interaction with human CD26. Emerg Microbes Infect. 2020;9(1):601–604.

23. Goyal P, Choi JJ, Pinheiro LC, et al. Clinical Characteristics of Covid-19 in New York City. N Engl J Med. 2020.

24. Wathelet MG, Orr M, Frieman MB, Baric RS. Severe acute respiratory syndrome coronavirus evades antiviral signaling: role of nsp1 and rational design of an attenuated strain. J Virol. 2007;81(21):11620–11633.

25. Tanaka Y, Sato Y, Sasaki T. Suppression of coronavirus replication by cyclophilin inhibitors. Viruses. 2013;5(5):1250–1260.

26. Zhao K, Li J, He W, et al. Cyclophilin B facilitates the replication of Orf virus. Virol J. 2017;14(1):114.

27. de Wilde AH, Zevenhoven-Dobbe JC, van der Meer Y, et al. Cyclosporin A inhibits the replication of diverse coronaviruses. J Gen Virol. 2011;92(Pt 11):2542–2548.

28. Pfefferle S, Schopf J, Kogl M, et al. The SARS-coronavirus-host interactome: identification of cyclophilins as target for pan-coronavirus inhibitors. PLoS Pathog. 2011;7(10):e1002331.

29. Pushkarsky T, Yurchenko V, Laborico A, Bukrinsky M. CD147 stimulates HIV-1 infection in a signal-independent fashion. Biochem Biophys Res Commun. 2007;363(3):495–499.

30. Kirk P, Wilson MC, Heddle C, Brown MH, Barclay AN, Halestrap AP. CD147 is tightly associated with lactate transporters MCT1 and MCT4 and facilitates their cell surface expression. EMBO J. 2000;19(15):3896–3904.

31. Ait-Ali N, Fridlich R, Millet-Puel G, et al. Rod-derived cone viability factor promotes cone survival by stimulating aerobic glycolysis. Cell. 2015;161(4):817–832.

32. Slomiany MG, Grass GD, Robertson AD, et al. Hyaluronan, CD44, and emmprin regulate lactate efflux and membrane localization of monocarboxylate transporters in human breast carcinoma cells. Cancer Res. 2009;69(4):1293–1301.

33. Berditchevski F, Chang S, Bodorova J, Hemler ME. Generation of monoclonal antibodies to integrin-associated proteins. Evidence that alpha3beta1 complexes with EMMPRIN/basigin/OX47/M6. J Biol Chem. 1997;272(46):29174–29180.

34. Xu D, Hemler ME. Metabolic activation-related CD147-CD98 complex. Mol Cell Proteomics. 2005;4(8):1061–1071.

35. Khunkaewla P, Schiller HB, Paster W, et al. LFA-1-mediated leukocyte adhesion regulated by interaction of CD43 with LFA-1 and CD147. Mol Immunol. 2008;45(6):1703–1711.

36. Priglinger CS, Szober CM, Priglinger SG, et al. Galectin-3 induces clustering of CD147 and integrin-beta1 transmembrane glycoprotein receptors on the RPE cell surface. PLoS One. 2013;8(7):e70011.

37. Till A, Rosenstiel P, Brautigam K, et al. A role for membrane-bound CD147 in NOD2-mediated recognition of bacterial cytoinvasion. J Cell Sci. 2008;121(Pt 4):487–495.

38. Zhou S, Zhou H, Walian PJ, Jap BK. CD147 is a regulatory subunit of the gamma-secretase complex in Alzheimer’s disease amyloid beta-peptide production. Proc Natl Acad Sci U S A. 2005;102(21):7499–7504.

39. Yao H, Teng Y, Sun Q, et al. Important functional roles of basigin in thymocyte development and T cell activation. Int J BiolSci. 2013;10(1):43–52.

40. Ruiz S, Castro-Castro A, Bustelo XR. CD147 inhibits the nuclear factor of activated T-cells by impairing Vav1 and Rac1 downstream signaling. J Biol Chem. 2008;283(9):5554–5566.

41. Muller MR, Rao A. NFAT, immunity and cancer: a transcription factor comes of age. Nat Rev Immunol. 2010;10(9):645–656.

42. Macian F. NFAT proteins: key regulators of T-cell development and function. Nat Rev Immunol. 2005;5(6):472–484.

43. Solstad T, Bains SJ, Landskron J, et al. CD147 (Basigin/Emmprin) identifies FoxP3+CD45RO+CTLA4+-activated human regulatory T cells. Blood. 2011;118(19):5141–5151.

44. Kim J, Yang YL, Jeong Y, Jang YS. Middle East Respiratory Syndrome-Coronavirus Infection into Established hDPP4-Transgenic Mice Accelerates Lung Damage Via Activation of the Pro-Inflammatory Response and Pulmonary Fibrosis. J Microbiol Biotechnol. 2020;30(3):427–438.

45. van Doremalen N, Miazgowicz KL, Milne-Price S, et al. Host species restriction of Middle East respiratory syndrome coronavirus through its receptor, dipeptidyl peptidase 4. J Virol. 2014;88(16):9220–9232.

46. Kleine-Weber H, Schroeder S, Kruger N, et al. Polymorphisms in dipeptidyl peptidase 4 reduce host cell entry of Middle East respiratory syndrome coronavirus. Emerg Microbes Infect. 2020;9(1):155–168.

47. Robinson MD, McCarthy DJ, Smyth GK. edgeR: a Bioconductor package for differential expression analysis of digital gene expression data. Bioinformatics. 2010;26(1):139–140.

48. Zou X, Chen K, Zou J, Han P, Hao J, Han Z. Single-cell RNA-seq data analysis on the receptor ACE2 expression reveals the potential risk of different human organs vulnerable to 2019-nCoV infection. Front Med. 2020.

49. Uhlen M, Fagerberg L, Hallstrom BM, et al. Proteomics. Tissue-based map of the human proteome. Science. 2015;347(6220):1260419.

50. Lu X, Zhang L, Du H, et al. SARS-CoV-2 Infection in Children. N Engl J Med. 2020;382(17):1663–1665.

51. Wu Z, McGoogan JM. Characteristics of and Important Lessons From the Coronavirus Disease 2019 (COVID-19) Outbreak in China: Summary of a Report of 72314 Cases From the Chinese Center for Disease Control and Prevention. JAMA. 2020.

52. Erkeller-Yuksel FM, Deneys V, Yuksel B, et al. Age-related changes in human blood lymphocyte subpopulations. J Pediatr. 1992;120(2 Pt 1):216–222.

53. Watanabe M, Risi R, Tuccinardi D, Baquero CJ, Manfrini S, Gnessi L. Obesity and SARS-CoV-2: a population to safeguard. Diabetes Metab Res Rev. 2020:e3325.

54. Recalcati S. Cutaneous manifestations in COVID-19: a first perspective. J Eur Acad Dermatol Venereol. 2020.

55. Kursat Azkur A, Akdis M, Azkur D, et al. Immune response to SARS-CoV-2 and mechanisms of immunopathological changes in COVID-19. Allergy. 2020;In Press.

56. Wang Q, Zhang Y, Wu L, et al. Structural and Functional Basis of SARS-CoV-2 Entry by Using Human ACE2. Cell. 2020.

57. Sungnak W, Huang N, Becavin C, et al. SARS-CoV-2 entry factors are highly expressed in nasal epithelial cells together with innate immune genes. Nat Med. 2020.

58. Ziegler CGK, Allon SJ, Nyquist SK, et al. SARS-CoV-2 receptor ACE2 is an interferon-stimulated gene in human airway epithelial cells and is detected in specific cell subsets across tissues. Cell. 2020.

59. Bröer S. The role of the neutral amino acid transporter B0AT1 (SLC6A19) in Hartnup disorder and protein nutrition. IUBMB life. 2009;61(6):591–599.

60. Leung JM, Yang CX, Tam A, et al. ACE-2 Expression in the Small Airway Epithelia of Smokers and COPD Patients: Implications for COVID-19. European Respiratory Journal. 2020:2000688.

61. Leung JM, Yang CX, Tam A, et al. ACE-2 Expression in the Small Airway Epithelia of Smokers and COPD Patients: Implications for COVID-19. Eur Respir J. 2020.

62. Imai Y, Kuba K, Rao S, et al. Angiotensin-converting enzyme 2 protects from severe acute lung failure. Nature. 2005;436(7047):112–116.

63. Goyal P, Choi JJ, Pinheiro LC, et al. Clinical Characteristics of Covid-19 in New York City. New England Journal of Medicine. 2020.

64. Jackson DJ, Busse WW, Bacharier LB, et al. Association of Respiratory Allergy, Asthma and Expression of the SARS-CoV-2 Receptor, ACE2. J Allergy Clin Immunol. 2020.

65. Peng L, Liu J, Xu W, et al. SARS-CoV-2 can be detected in urine, blood, anal swabs and oropharyngeal swabs specimens. J Med Virol. 2020.

66. Bao W, Min D, Twigg SM, et al. Monocyte CD147 is induced by advanced glycation end products and high glucose concentration: possible role in diabetic complications. Am J Physiol Cell Physiol. 2010;299(5):C1212–1219.

67. Zhang H, Fan Q, Xie H, et al. Elevated Serum Cyclophilin B Levels Are Associated with the Prevalence and Severity of Metabolic Syndrome. Frontiers in Endocrinology. 2017;8(360).

68. Hahn JN, Kaushik DK, Yong VW. The role of EMMPRIN in T cell biology and immunological diseases. J Leukoc Biol. 2015;98(1):33–48.

69. Halestrap AP. The monocarboxylate transporter family--Structure and functional characterization. IUBMB Life. 2012;64(1):1–9.

70. Darmoul D, Voisin T, Couvineau A, et al. Regional expression of epithelial dipeptidyl peptidase IV in the human intestines. Biochem Biophys Res Commun. 1994;203(2):1224–1229.

71. Li Y, Jiang D, Liang J, et al. Severe lung fibrosis requires an invasive fibroblast phenotype regulated by hyaluronan and CD44. J Exp Med. 2011;208(7):1459–1471.

72. Pure E, Cuff CA. A crucial role for CD44 in inflammation. Trends Mol Med. 2001;7(5):213–221.

73. Chen H, Hou J, Jiang X, et al. Response of memory CD8+ T cells to severe acute respiratory syndrome (SARS) coronavirus in recovered SARS patients and healthy individuals. Journal of immunology (Baltimore, Md: 1950). 2005;175(1):591–598.

74. Bouaziz JD, Duong T, Jachiet M, et al. Vascular skin symptoms in COVID-19: a french observational study. J Eur Acad Dermatol Venereol. 2020.

75. Wollenberg A, Flohr C, Simon D, et al. European Task Force on Atopic Dermatitis (ETFAD) statement on severe acute respiratory syndrome coronavirus 2 (SARS-Cov-2)-infection and atopic dermatitis. J Eur Acad Dermatol Venereol. 2020.

76. Seegraber M, Worm M, Werfel T, et al. Recurrent eczema herpeticum -a retrospective European multicenter study evaluating the clinical characteristics of eczema herpeticum cases in atopic dermatitis patients. J Eur Acad Dermatol Venereol. 2019.

77. Vankadari N, Wilce JA. Emerging COVID-19 coronavirus: glycan shield and structure prediction of spike glycoprotein and its interaction with human CD26. Emerging Microbes & Infections. 2020;9(1):601–604.

78. Allan DSJ, Cerdeira AS, Ranjan A, et al. Transcriptome analysis reveals similarities between human blood CD3(-) CD56(bright) cells and mouse CD127(+) innate lymphoid cells. Sci Rep. 2017;7(1):3501.

79. Peters MJ, Joehanes R, Pilling LC, et al. The transcriptional landscape of age in human peripheral blood. Nat Commun. 2015;6:8570.

80. Team CC-R. Coronavirus Disease 2019 in Children -United States, February 12-April 2, 2020. MMWR Morb Mortal Wkly Rep. 2020;69(14):422–426.

