## Supplementary Figures for "Distribution of ACE2, CD147, cyclophilins, CD26 and other SARS-CoV-2 associated molecules in human tissues and immune cells in health and disease"

**Supplementary Figure Titles and Legends**

**Figure S1. Heatmap of ACE-2-, CD147-and CD26-related gene expression in different cells and tissues**

**Figure S2. Gene Co-Expression in Bronchial Biopsy in non-diseased control**

**Figure S3. Gene Co-Expression in Blood in non-diseased control**

**Figure S4. Human Bronchial Epithelial Cells**

**A-E** Gene expression of ACE, SDC1， LGALS3, NFAT5 and NME1 in control, asthma and COPD population

**Figure S5. Bronchial Biopsy**

**A-F** Gene expression of ACE, ACE2, BSG(CD147), CD44, SLC7A5(CD98) and ITGA3 in subgroups including control, asthma, COPD, non-obese, obese, normotension, hypertension, non-smoker, smoker, female and male.

**G-H** Gene expression of ITGA6 and APOD in subgroups including control, asthma, COPD, non-obese, obese, normotension, hypertension, non-smoker, smoker, female and male.

**Figure S6. Gene Co-Expression in Bronchial Biopsy in asthma patients**

**Figure S7. Broncho-Alveolar Lavage**

**A-F** Gene expression of ACE, ACE2, BSG(CD147), PPIA, S100A9 and CD44 in subgroups including control, asthma, COPD, non-obese, obese, normotension, hypertension, non-smoker, smoker, female and male.

**G-L** Gene expression of SLC16A7(MCT2), SLC16A3(MCTs), ITGA3, NFATC1, NFATC2 and SLC7A5(CD98) in subgroups including control, asthma, COPD, non-obese, obese, normotension, hypertension, non-smoker, smoker, female and male.

**M-O** Gene expression of APH1A, PSEN1 and PSENEN in subgroups including control, asthma, COPD, non-obese, obese, normotension, hypertension, non-smoker, smoker, female and male.

**Figure S8. Blood**

**A-F** Gene expression of ACE, ACE2, BSG(CD147), PPIA, S100A9 and CD44 in subgroups including control, asthma, COPD, non-obese, obese, normotension, hypertension, non-smoker, smoker, female and male.

**G-L** Gene expression of SLC16A7(MCT2), ITGA6, NFATC2, LGALS3, NOD2 and NME1 in subgroups including control, asthma, COPD, non-obese, obese, normotension, hypertension, non-smoker, smoker, female and male.

**M** Gene expression of DPP4 in subgroups including control, asthma, COPD, non-obese, obese, normotension, hypertension, non-smoker, smoker, female and male.

**Figure S9. Gene Co-Expression in Broncho-Alveolar Lavage in asthma patients**

**Figure S10. Gene Co-Expression in Blood in asthma patients**

**Figure S11. Correlation Analysis**

**A** Correlation analysis between SDC1 and age in broncho-alveolar lavage

**B** Correlation analysis between SLC16A1 and age in blood

**C** Correlation analysis between NME1 and BMI in blood

**D** Correlation analysis between NCSTN and BMI in blood

### Figure S1

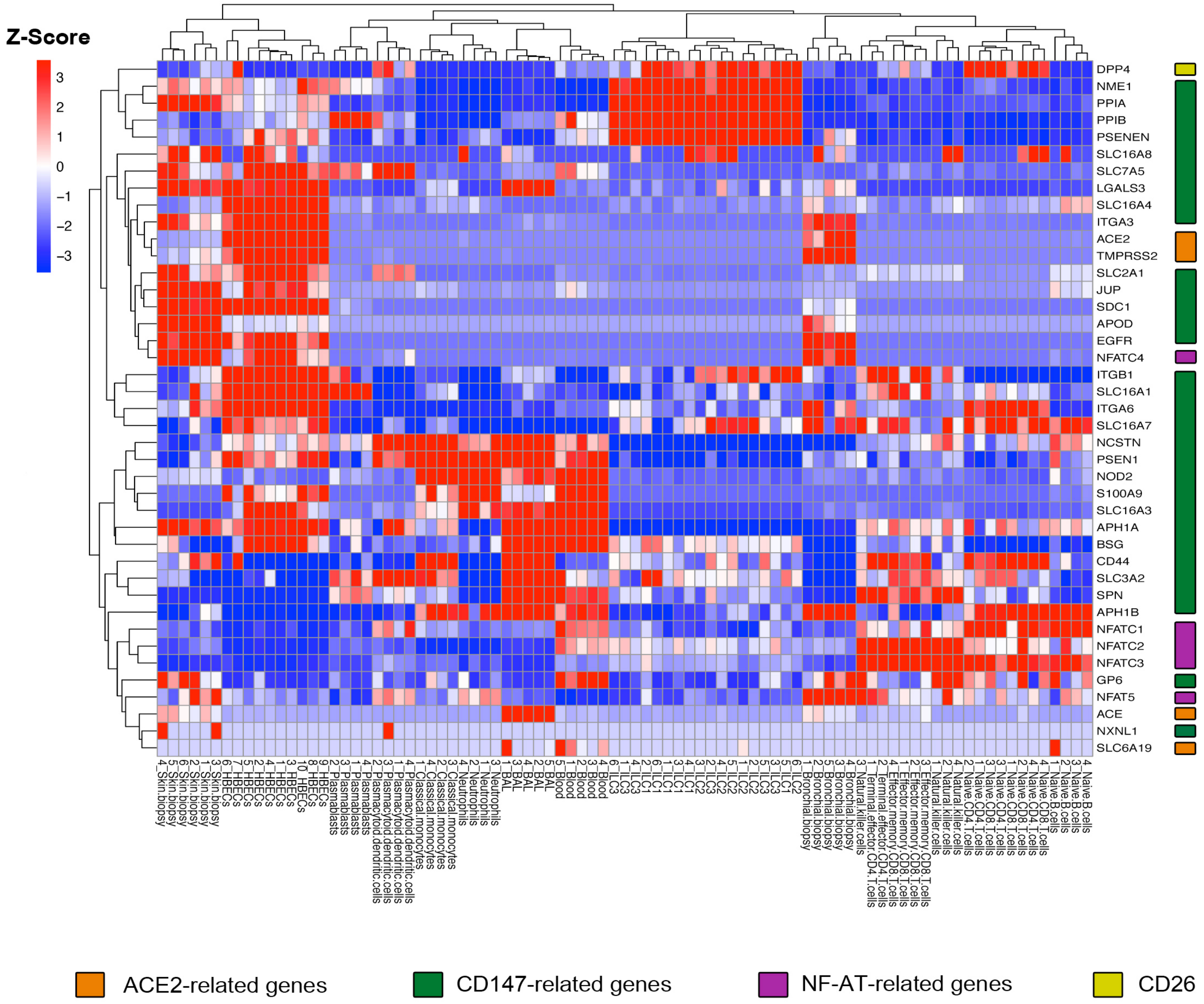
Z-Score
3
2
1
0
−1
−2
−3
DPP4
NME1
PPIA
PPIB
PSENEN
SLC16A8
SLC7A5
LGALS3
SLC16A4
ITGA3
ACE2
TMPRSS2
SLC2A1
JUP
SDC1
APOD
EGFR
NFATC4
ITGB1
SLC16A1
ITGA6
SLC16A7
NCSTN
PSEN1
NOD2
S100A9
SLC16A3
APH1A
BSG
CD44
SLC3A2
SPN
APH1B
NFATC1
NFATC2
NFATC3
GP6
NFAT5
ACE
NXNL1
SLC6A19
ACE2-related genes
CD147-related genes
NF-AT-related genes
CD26

### Figure S2

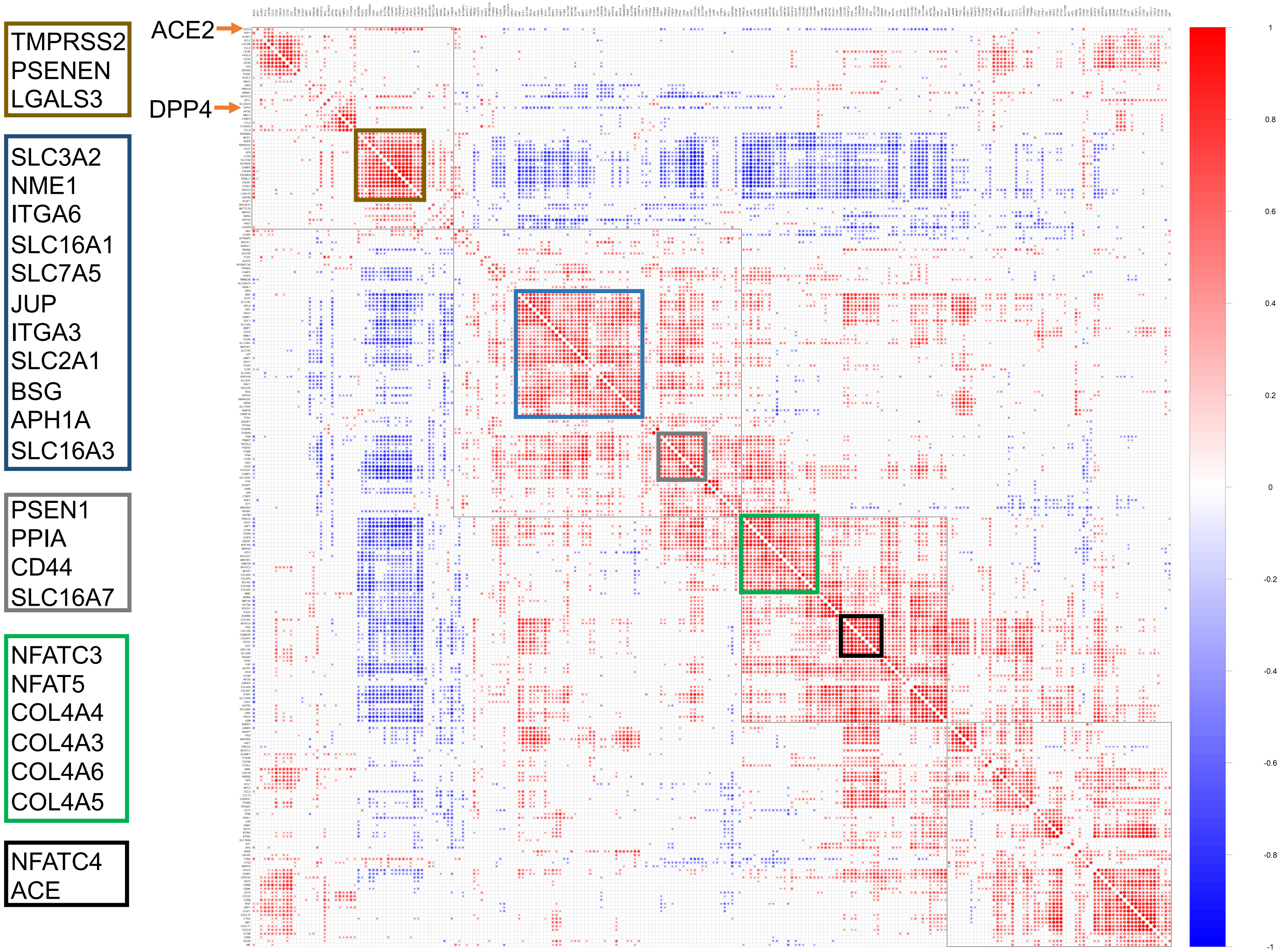
TMPRSS2
PSENEN
LGALS3
SLC3A2
NME1
ITGA6
SLC16A1
SLC7A5
JUP
ITGA3
SLC2A1
BSG
APH1A
SLC16A3
PSEN1
PPIA
CD44
SLC16A7
NFATC3
NFAT5
COL4A4
COL4A3
COL4A6
COL4A5
NFATC4
ACE
ACE2
DPP4
1
0.8
0.6
0.4
0.2
0
-0.2
-0.4
-0.6
-0.8
-1

### Figure S3

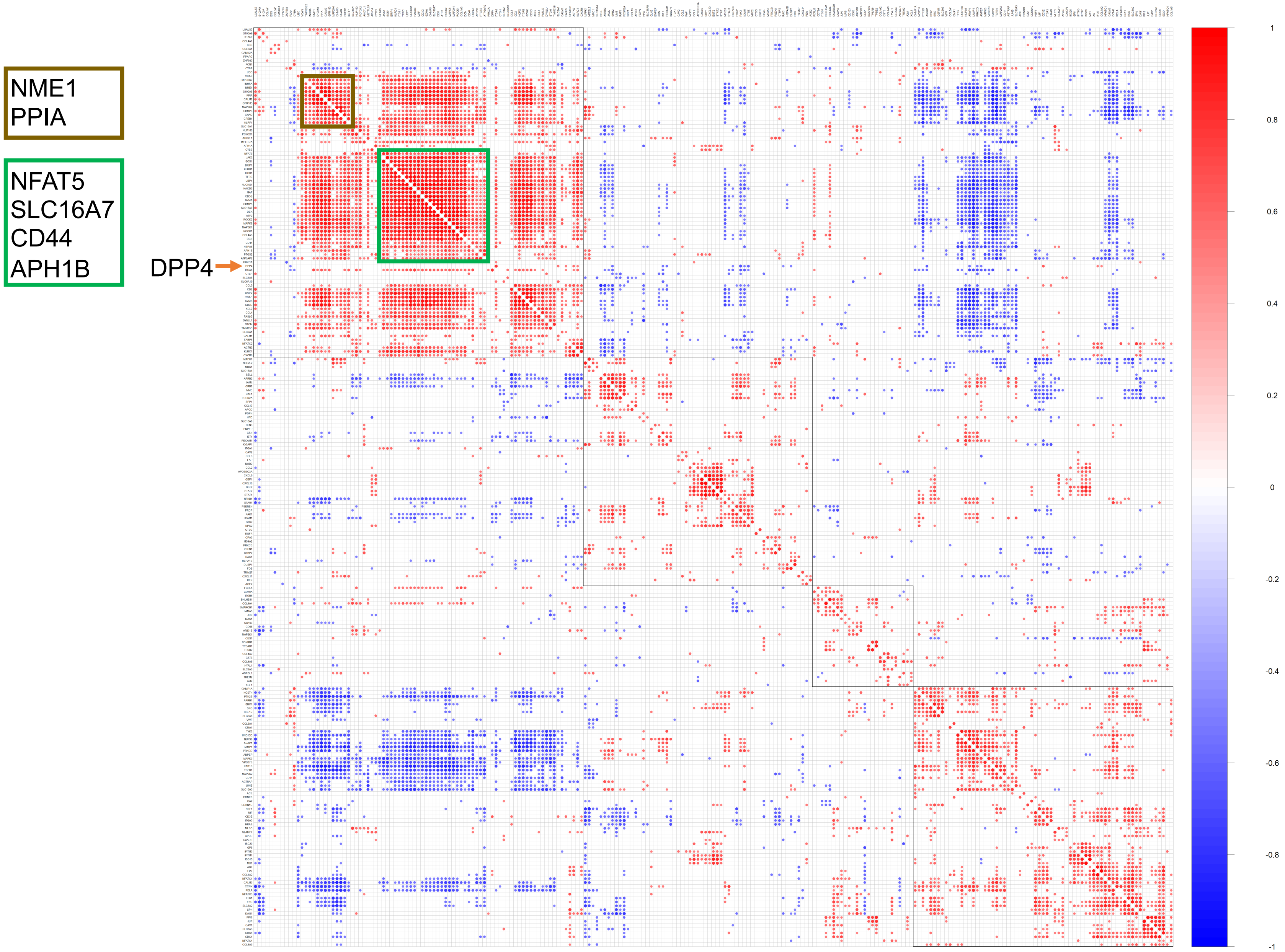
NME1
PPIA
NFAT5
SLC16A7
CD44
APH1B
DPP4
1
0.8
0.6
0.4
0.2
0
-0.2
-0.4
-0.6
-0.8
-1

### Figure S4

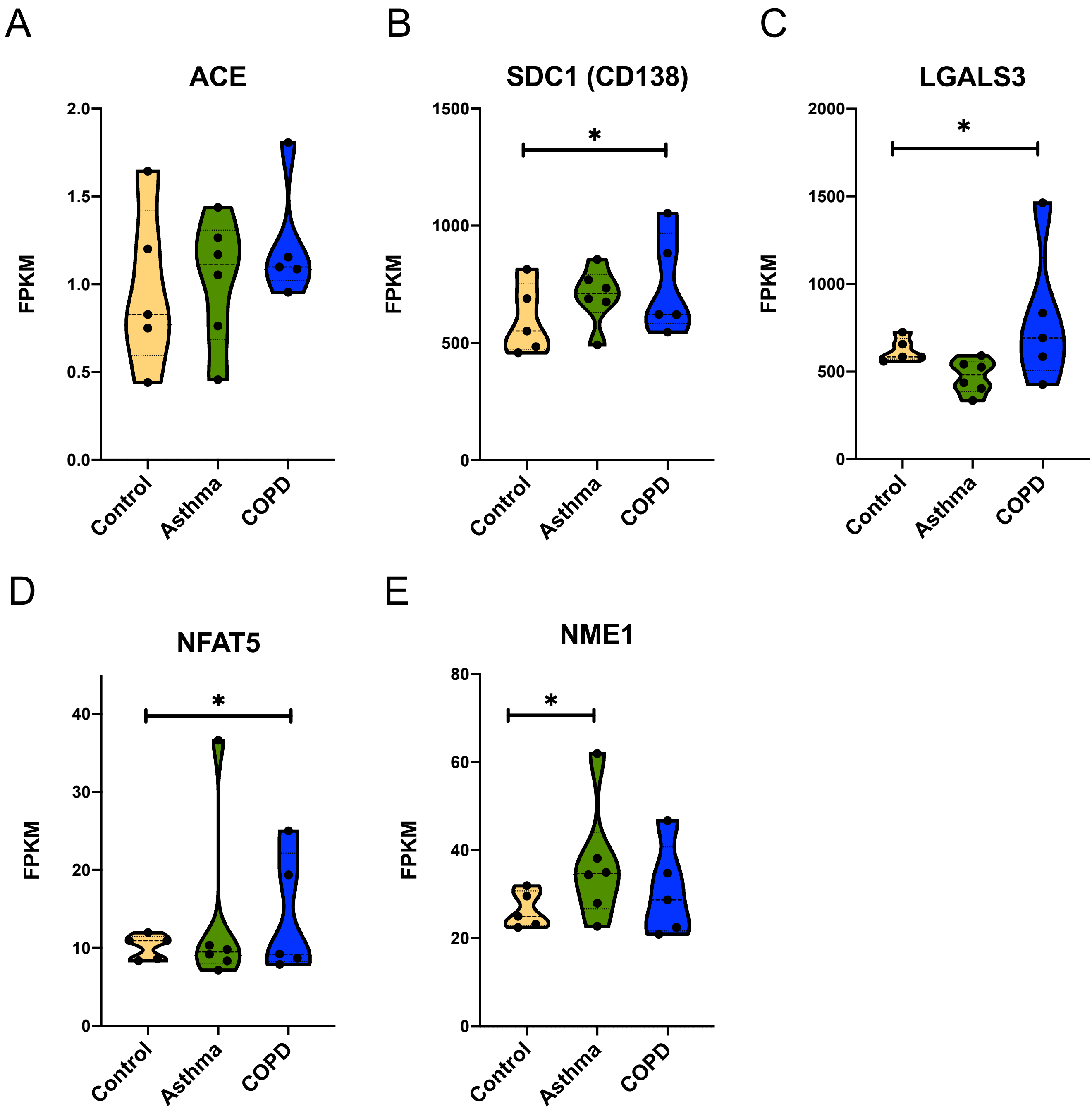
A
ACE
FPKM
2.0
1.5
1.0
0.5
0.0
Control
Asthma
COPD
B
SDC1 (CD138)
1500
1000
500
0
*
C
LGALS3
2000
1500
1000
500
0
*
D
NFAT5
40
30
20
10
0
*
E
NME1
80
60
40
20
0
*

### Figure S5

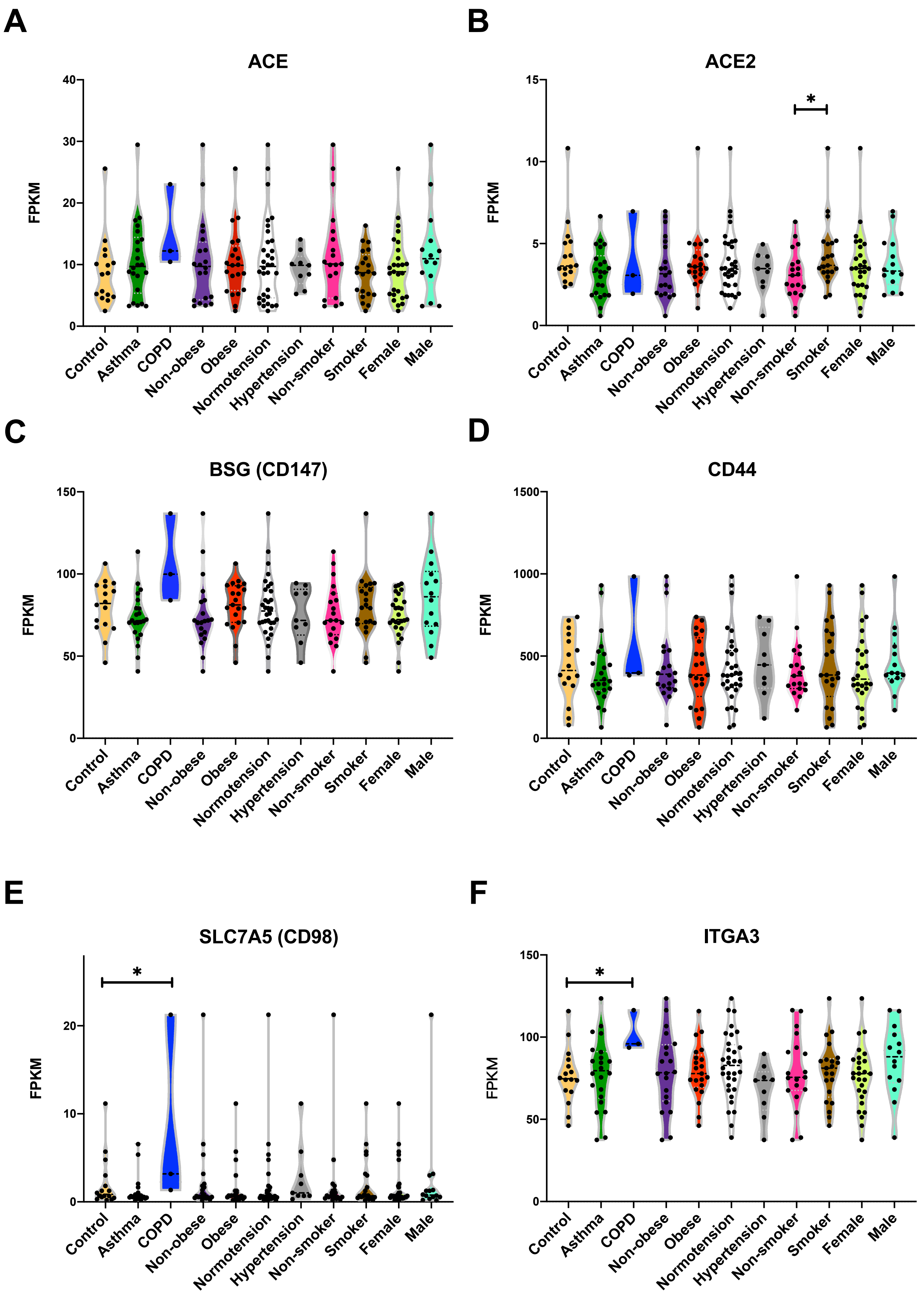
A
ACE
FPKM
40
30
20
10
0
Control
Asthma
COPD
Non-obese
Obese
Normotension
Hypertension
Non-smoker
Smoker
Female
Male
B
ACE2
FPKM
15
10
5
0
*
C
BSG (CD147)
FPKM
150
100
50
0
D
CD44
FPKM
1500
1000
500
0
E
SLC7A5 (CD98)
FPKM
20
10
0
*
F
ITGA3
FPKM
150
100
50
0
*

### Figure S5

G

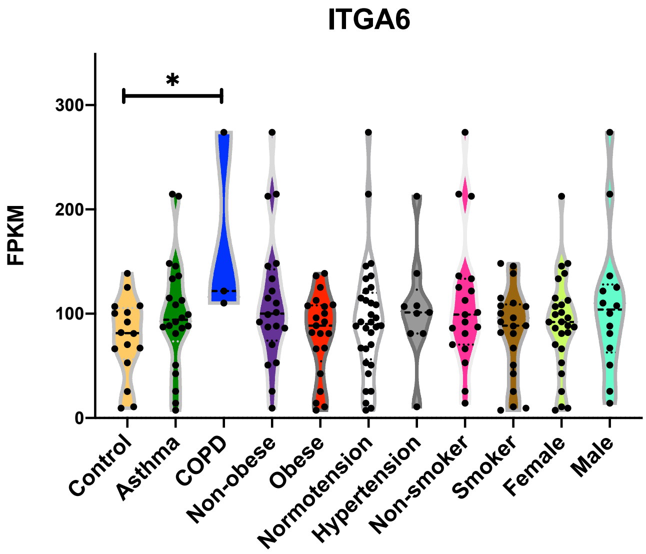
ITGA6
FPKM
300
200
100
0
*
Control
Asthma
COPD
Non-obese
Obese
Normotension
Hypertension
Non-smoker
Smoker
Female
Male

H

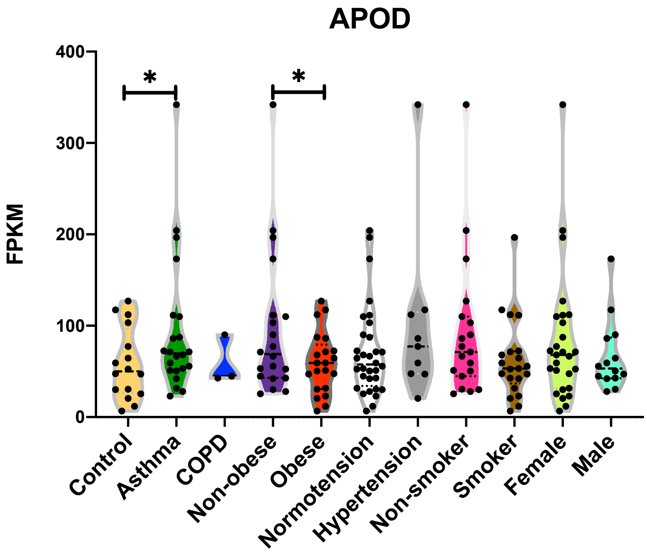
APOD
FPKM
400
300
200
100
0
*
*
Control
Asthma
COPD
Non-obese
Obese
Normotension
Hypertension
Non-smoker
Smoker
Female
Male

### Figure S6

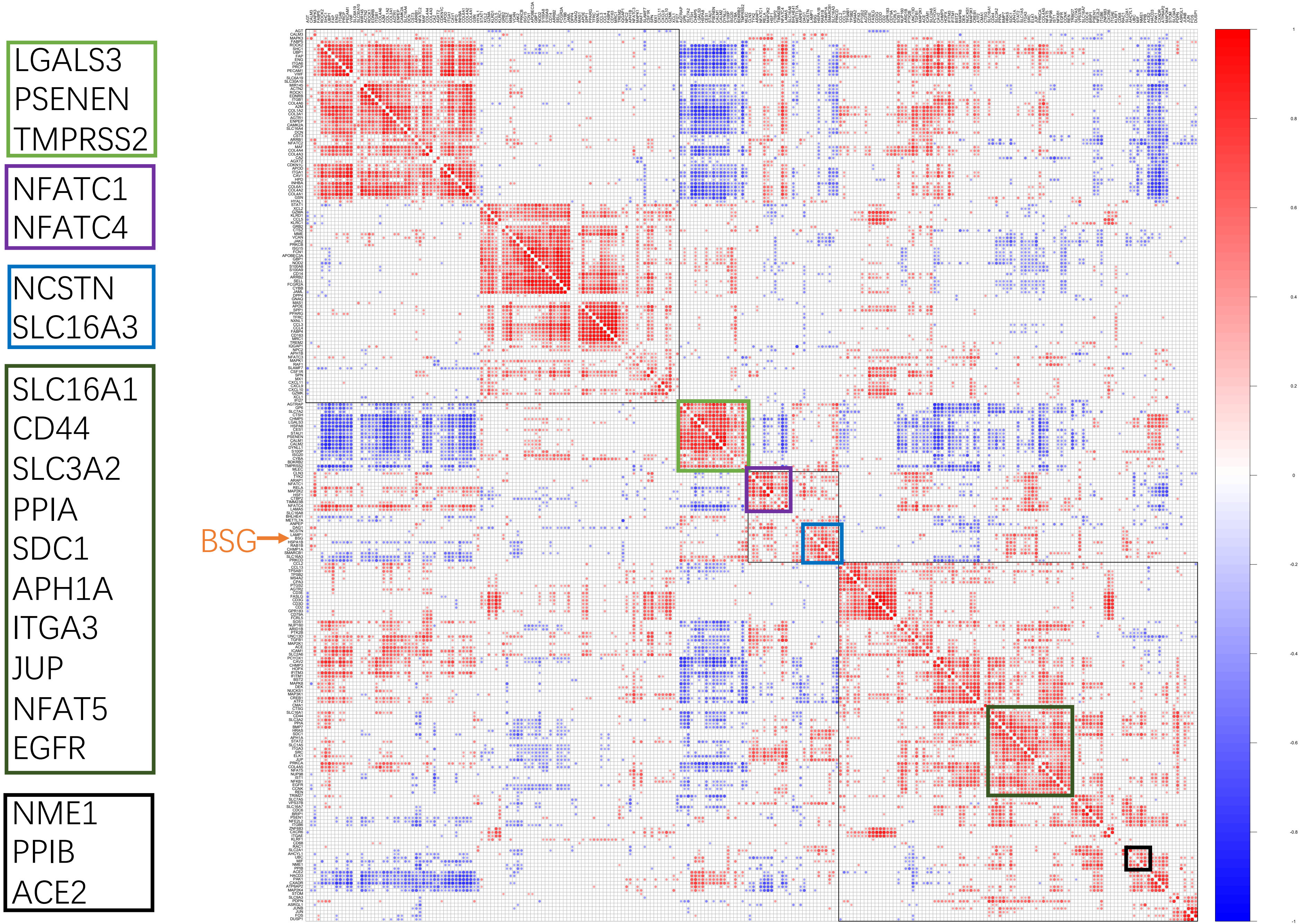
LGALS3
PSENEN
TMPRSS2
NFATC1
NFATC4
NCSTN
SLC16A3
SLC16A1
CD44
SLC3A2
PPIA
SDC1
APH1A
ITGA3
JUP
NFAT5
EGFR
NME1
PPIB
ACE2
BSG

### Figure S7

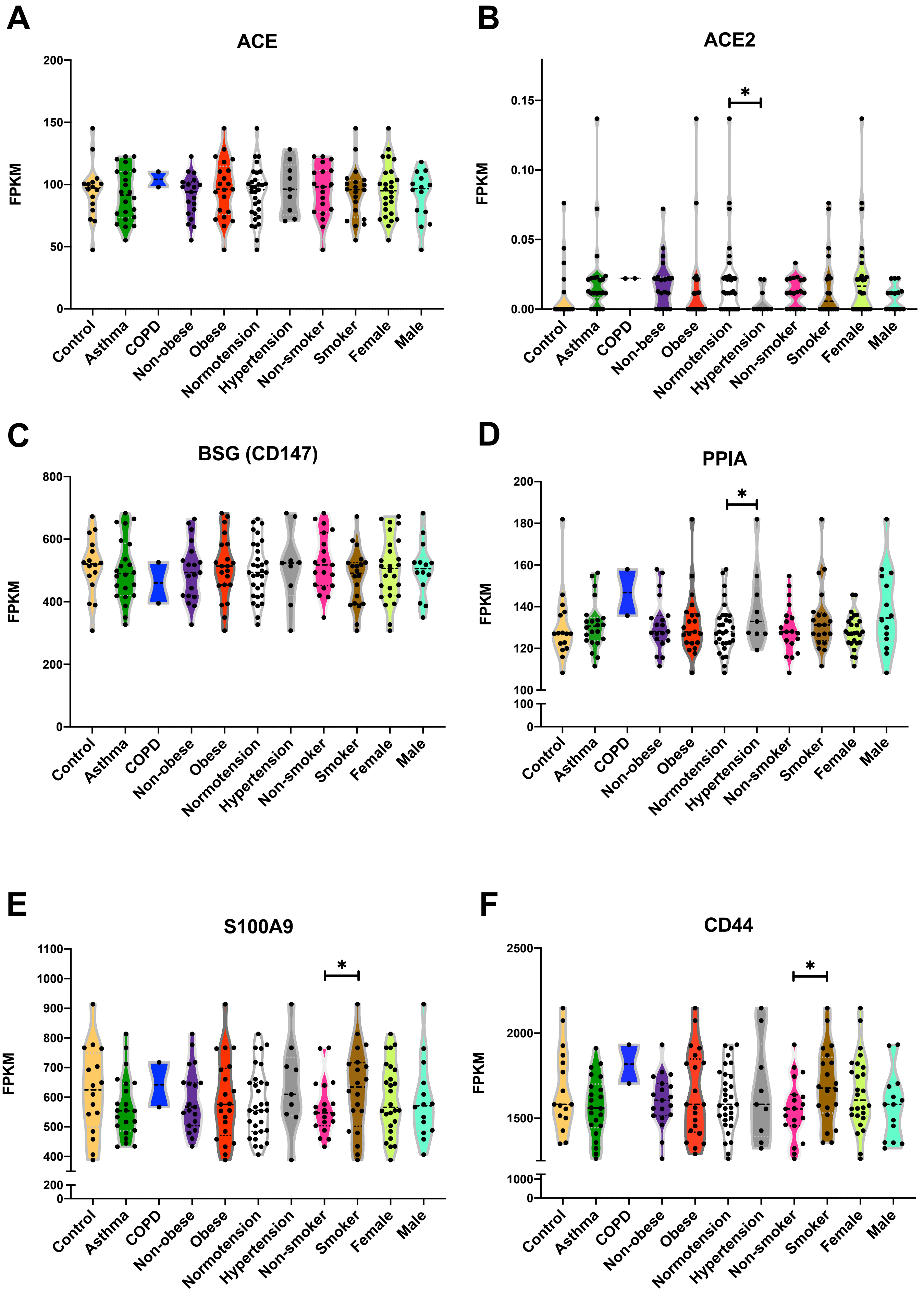
A
ACE
FPKM
B
ACE2
FPKM
*
C
BSG (CD147)
FPKM
D
PPIA
FPKM
*
E
S100A9
FPKM
*
F
CD44
FPKM
*
Control
Asthma
COPD
Non-obese
Obese
Normotension
Hypertension
Non-smoker
Smoker
Female
Male

### Figure S7

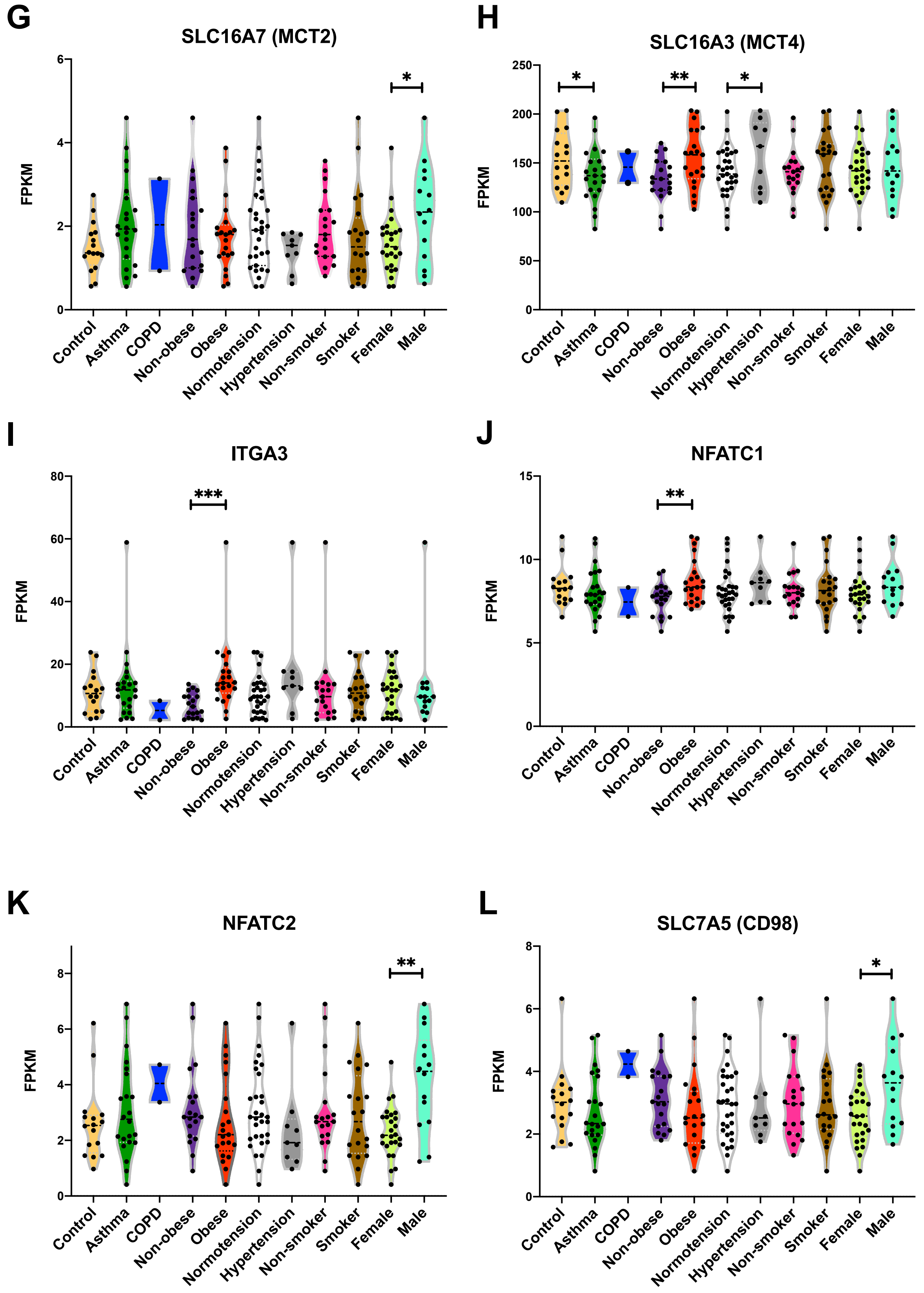
G
SLC16A7 (MCT2)
FPKM
0
2
4
6
*
H
SLC16A3 (MCT4)
FPKM
0
50
100
150
200
250
*
**
*
I
ITGA3
FPKM
0
20
40
60
80
***
J
NFATC1
FPKM
0
5
10
15
**
K
NFATC2
FPKM
0
2
4
6
8
**
L
SLC7A5 (CD98)
FPKM
0
2
4
6
8
*
Control
Asthma
COPD
Non-obese
Obese
Normotension
Hypertension
Non-smoker
Smoker
Female
Male

### Figure S7

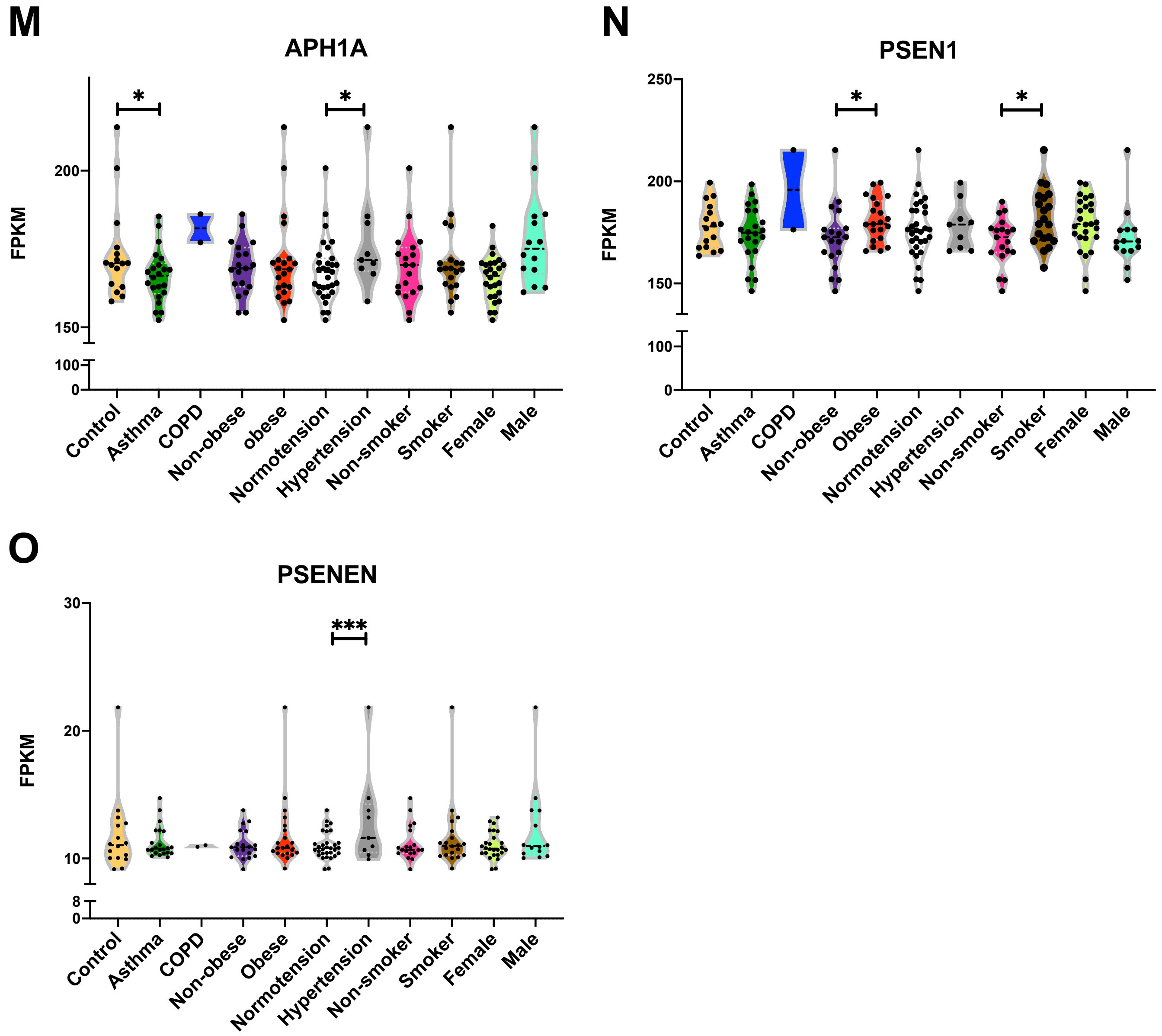
M
APH1A
FPKM
200
150
100
0
*
*
Control
Asthma
COPD
Non-obese
obese
Normotension
Hypertension
Non-smoker
Smoker
Female
Male
N
PSEN1
FPKM
250
200
150
100
0
*
*
Control
Asthma
COPD
Non-obese
Obese
Normotension
Hypertension
Non-smoker
Smoker
Female
Male
O
PSENEN
FPKM
30
20
10
8
0
***
Control
Asthma
COPD
Non-obese
Obese
Normotension
Hypertension
Non-smoker
Smoker
Female
Male

### Figure S8

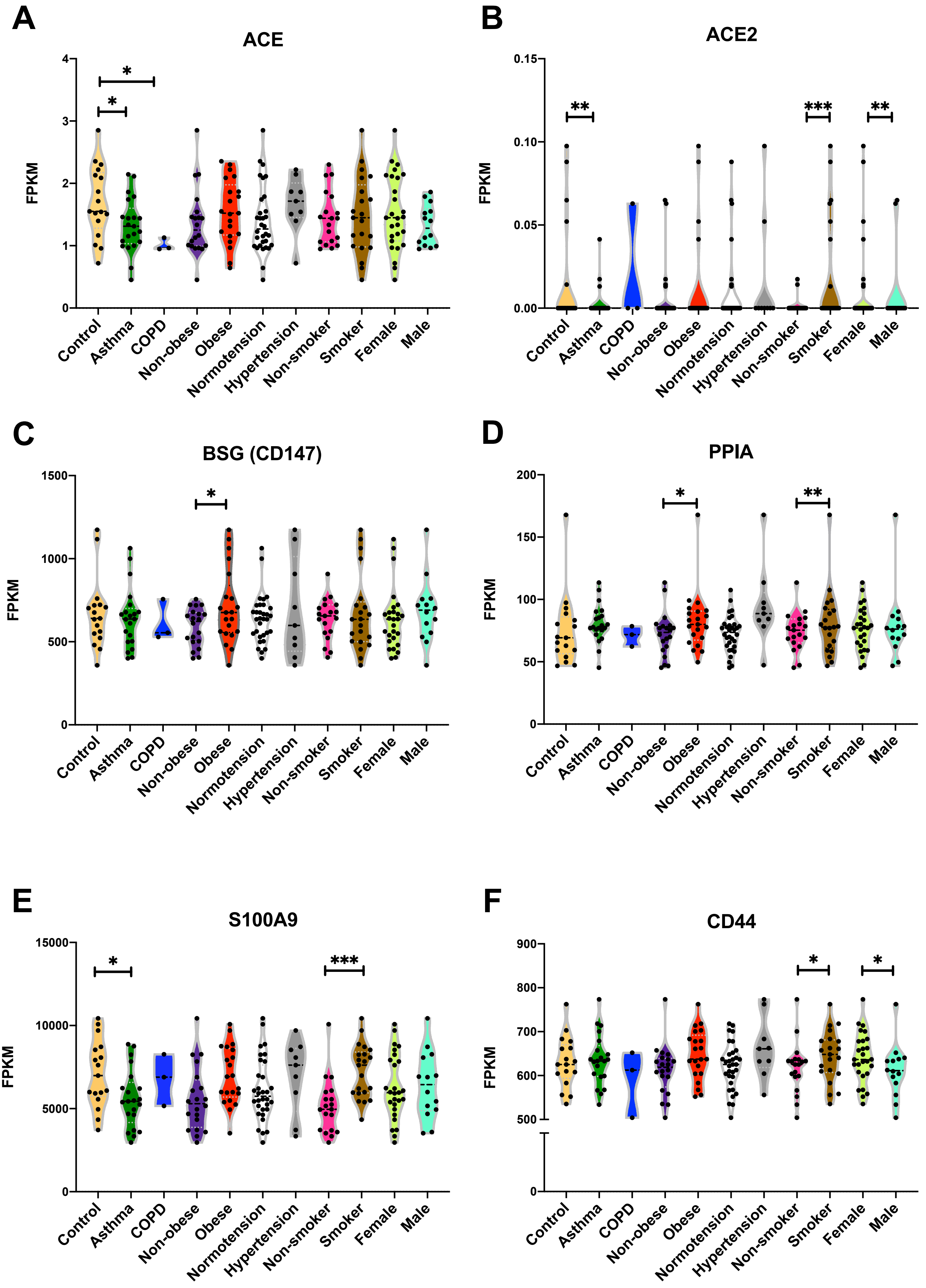
A
ACE
FPKM
0
1
2
3
4
*
*
B
ACE2
FPKM
0.00
0.05
0.10
0.15
**
***
**
C
BSG (CD147)
FPKM
0
500
1000
1500
*
D
PPIA
FPKM
0
50
100
150
200
*
**
E
S100A9
FPKM
0
5000
10000
15000
*
***
F
CD44
FPKM
0
500
600
700
800
900
*
*
Control
Asthma
COPD
Non-obese
Obese
Normotension
Hypertension
Non-smoker
Smoker
Female
Male

### Figure S8

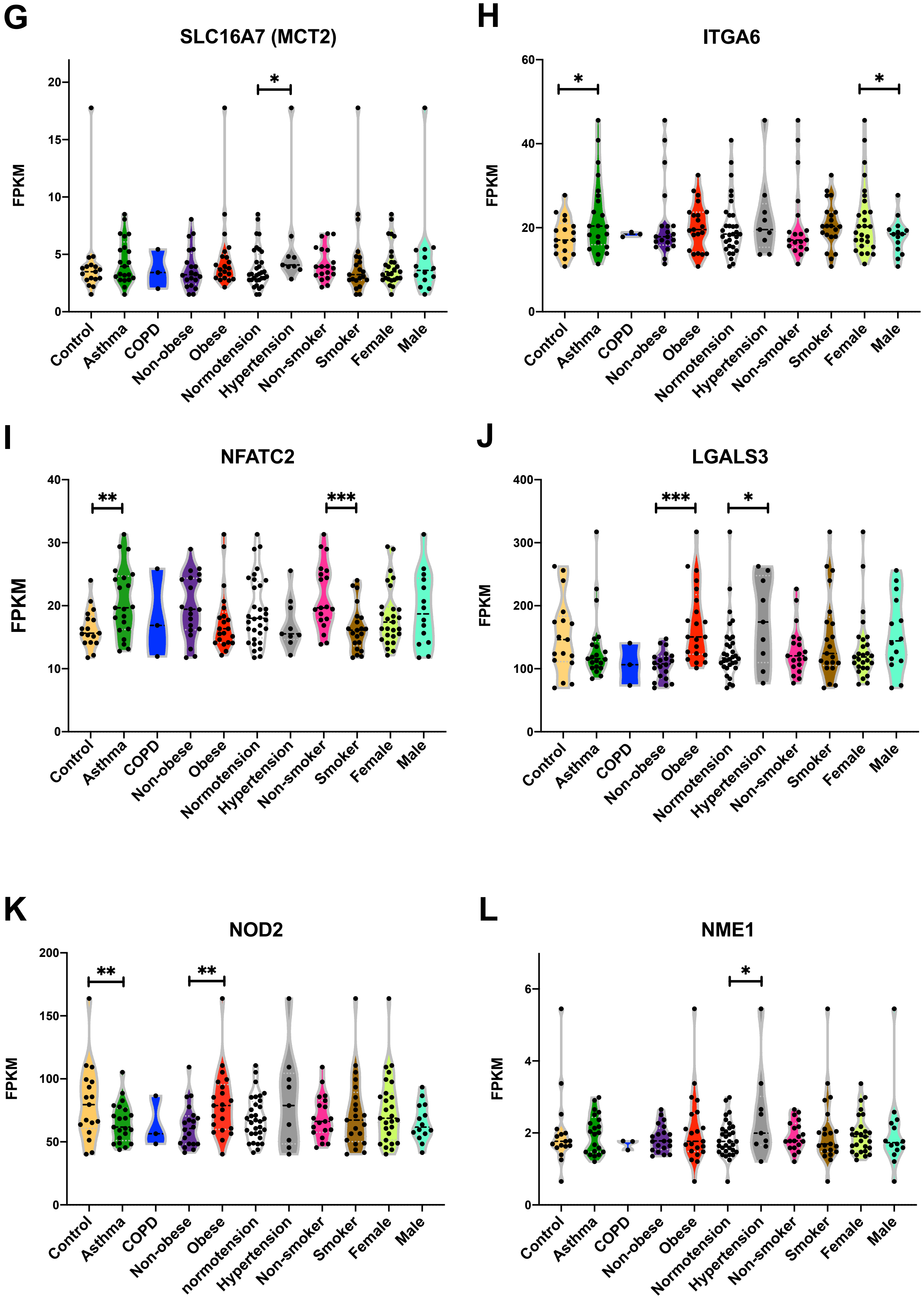
G
SLC16A7 (MCT2)
FPKM
0
5
10
15
20
*
Control
Asthma
COPD
Non-obese
Obese
Normotension
Hypertension
Non-smoker
Smoker
Female
Male
H
ITGA6
FPKM
0
20
40
60
*
*
Control
Asthma
COPD
Non-obese
Obese
Normotension
Hypertension
Non-smoker
Smoker
Female
Male
I
NFATC2
FPKM
0
10
20
30
40
**
***
Control
Asthma
COPD
Non-obese
Obese
Normotension
Hypertension
Non-smoker
Smoker
Female
Male
J
LGALS3
FPKM
0
100
200
300
400
***
*
Control
Asthma
COPD
Non-obese
Obese
Normotension
Hypertension
Non-smoker
Smoker
Female
Male
K
NOD2
FPKM
0
50
100
150
200
**
**
Control
Asthma
COPD
Non-obese
Obese
normotension
Hypertension
Non-smoker
Smoker
Female
Male
L
NME1
FPKM
0
2
4
6
*
Control
Asthma
COPD
Non-obese
Obese
Normotension
Hypertension
Non-smoker
Smoker
Female
Male

### Figure S8

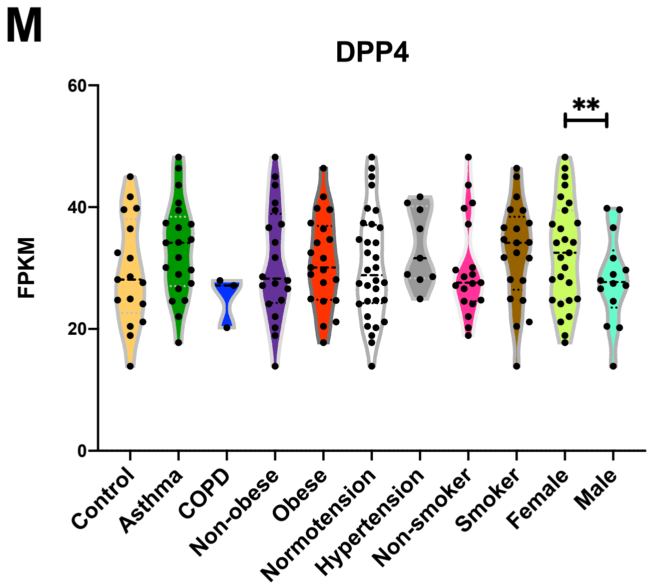
M
DPP4
60
40
20
0
FPKM
**
Control
Asthma
COPD
Non-obese
Obese
Normotension
Hypertension
Non-smoker
Smoker
Female
Male

#### Figure S9

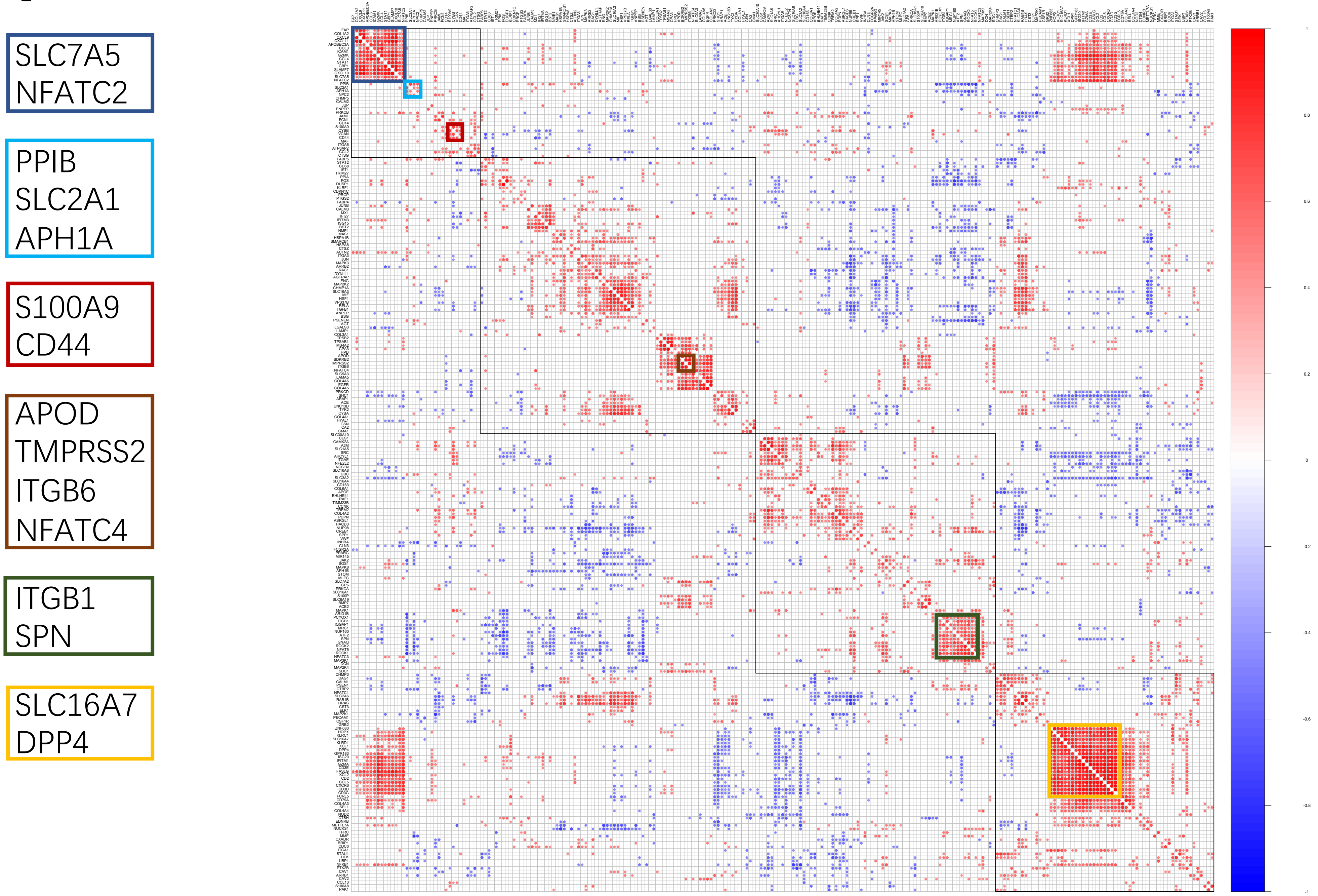
SLC7A5
NFATC2
PPIB
SLC2A1
APH1A
S100A9
CD44
APOD
TMPRSS2
ITGB6
NFATC4
ITGB1
SPN
SLC16A7
DPP4

### Figure S10

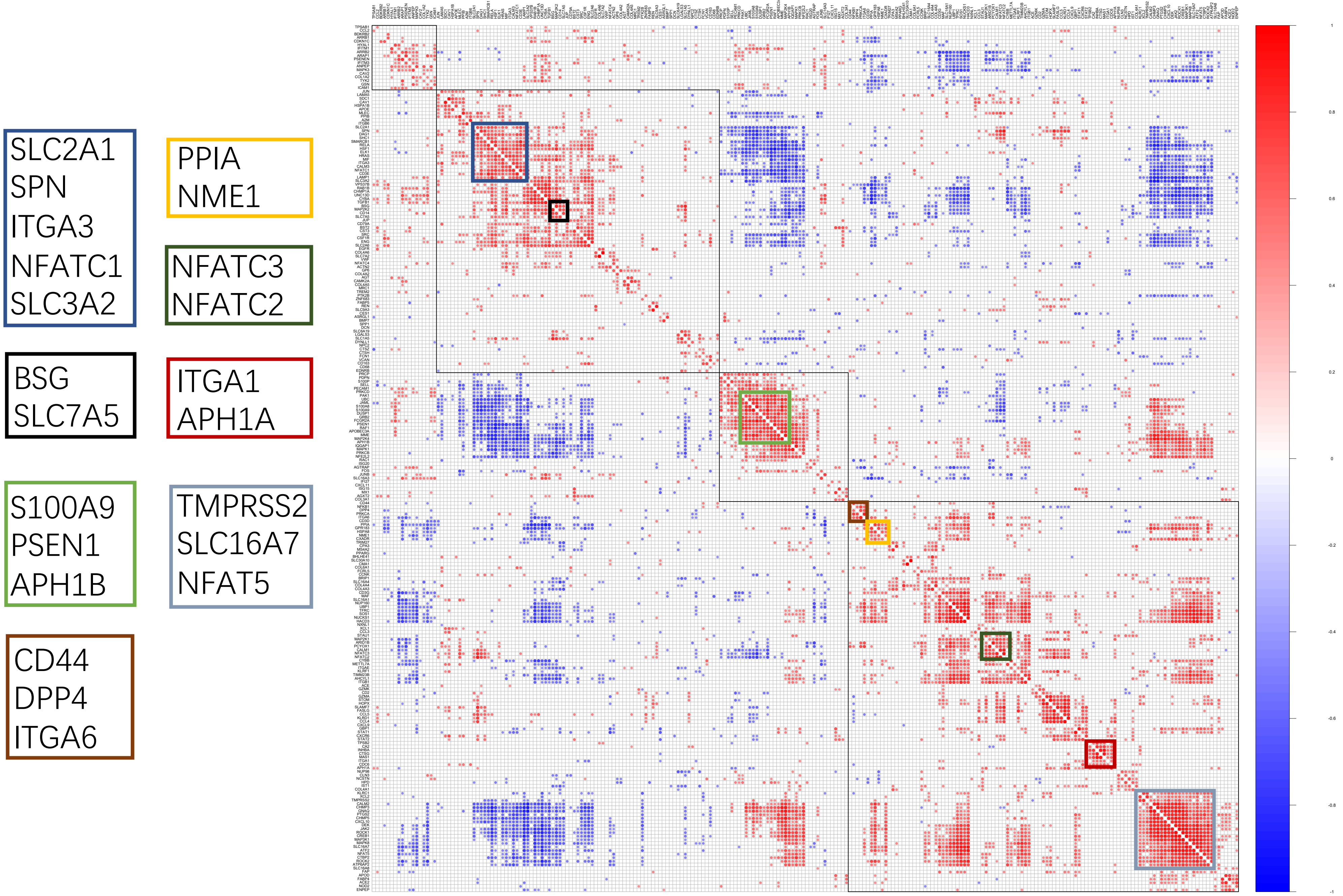

SLC2A1
SPN
ITGA3
NFATC1
SLC3A2
PPIA
NME1
NFATC3
NFATC2
BSG
SLC7A5
ITGA1
APH1A
S100A9
PSEN1
APH1B
TMPRSS2
SLC16A7
NFAT5
CD44
DPP4
ITGA6

### Figure S11

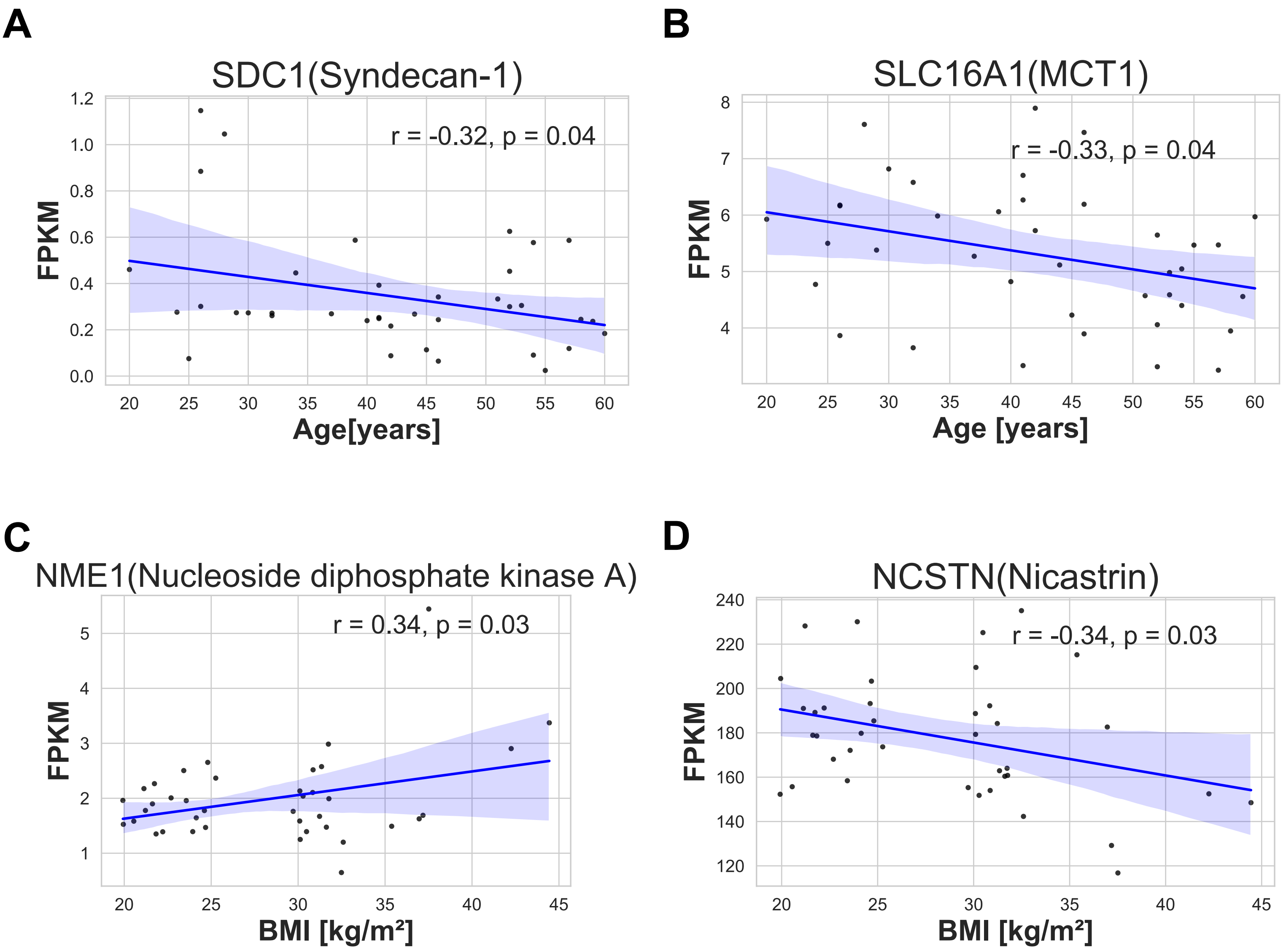
A
SDC1(Syndecan-1)
r = -0.32, p = 0.04
FPKM
1.2
1.0
0.8
0.6
0.4
0.2
0.0
20 25 30 35 40 45 50 55 60
Age[years]
B
SLC16A1(MCT1)
r = -0.33, p = 0.04
FPKM
8
7
6
5
4
20 25 30 35 40 45 50 55 60
Age [years]
C
NME1(Nucleoside diphosphate kinase A)
r = 0.34, p = 0.03
FPKM
5
4
3
2
1
20 25 30 35 40 45
BMI [kg/m²]
D
NCSTN(Nicastrin)
r = -0.34, p = 0.03
FPKM
240
220
200
180
160
140
120
20 25 30 35 40 45
BMI [kg/m²]
