## Supplementary Methods and Tables for "Distribution of ACE2, CD147, cyclophilins, CD26 and other SARS-CoV-2 associated molecules in human tissues and immune cells in health and disease"

**Short title:** SARS-CoV-2 associated molecules in health and disease

Radzikowska U.[1,2,3]*, Ding M.[1,2,4]*, Tan G.[1,5], Zhakparov D.[1], Peng Y.[1,2,6], Wawrzyniak P.[1,2,7], Wang M.[1,2,8], Li S.[1,9], Morita H.[1,10], Altunbulakli C.[1,2], Reiger M.[11], Neumann AU.[11,12,13], Lunjani N.[1,2], Traidl-Hoffmann C.[2,11], Nadeau K.[14], O'Mahony L.[1,15], Akdis CA.[1,2], Sokolowska M.[1,2#]

***equal contribution**

**Affiliations**

[1] Swiss Institute of Allergy and Asthma Research (SIAF), University of Zurich, Davos, Switzerland

[2] Christine Kühne – Center for Research and Education (CK-CARE), Davos, Switzerland

[3] Department of Regenerative Medicine and Immune Regulation, Medical University of Bialystok, Bialystok, Poland

[4] Department of Allergology, Zhongnan Hospital of Wuhan University, Wuhan, China

[5] Functional Genomic Centre Zurich, ETH Zurich/University of Zurich, Zurich, Switzerland

[6] Otorhinolaryngology Hospital, The First Affiliated Hospital, Sun Yat-sen University, Guangzhou, China

[7] Division of Clinical Chemistry and Biochemistry, University Children`s Hospital Zurich, Zurich, Switzerland; Children`s Research Center, University Children`s Hospital Zurich, Zurich, Switzerland

[8] Department of Otolaryngology, Head and Neck Surgery, Beijing TongRen Hospital, Capital Medical University and the Beijing Key Laboratory of Nasal Diseases, Beijing Institute of Otolaryngology, Beijing, China

[9] Department of Cancer Immunology, Institute for Cancer Research, Oslo University Hospital, Oslo, Norway

[10] Department of Allergy and Clinical Immunology, National Research Institute for Child Health and Development, Tokyo, Japan

[11] Chair and Institute of Environmental Medicine, UNIKA-T, Technical University of Munich and Helmholtz Zentrum Munchen, Augsburg, Germany

[12] Institute of Computational Biology (ICB), Helmholtz Zentrum Munchen, Munich, Germany

[13] Institute of Experimental Medicine (IEM), Czech Academy of Sciences, Prague, Czech Republic

[14] Sean N Parker Centre for Allergy and Asthma Research at Stanford University, Department of Medicine, Stanford University School of Medicine, Stanford, USA

[15] Department of Medicine and School of Microbiology, APC Microbiome Ireland, National University of Ireland, Cork, Ireland

**# corresponding author:**

Milena Sokolowska MD, PhD
Head Immune Metabolism
Swiss Institute of Allergy and Asthma Research (SIAF)
University of Zurich
Herman-Burchard-Strasse 9
CH-7265 Davos-Wolfgang

https://www.siaf.uzh.ch/immune_metabolism.html

### Online Supplementary

### Material and Methods

We analyzed gene expression of SARS-CoV-2 receptors and related molecules' (Table S1) in a broad range of tissues and immune cells from the human RNA-seq databases generated by our *ex vivo* and *in vitro* approaches in the Swiss Institute of Asthma and Allergy Research (SIAF), by our collaborators or from the Gene Expression Omnibus (http://www.ncbi.nlm.nih.gov/). Detailed description of projects is presented below:

#### SIAF Study 1

Differentiated primary human bronchial epithelial cells (HBECs) from controls, asthmatics, and COPD patients were processed as described in the original paper [1]. Briefly, total RNA was extracted and purified. Sequencing was done on the Illumina HiSeq 4000.

**SIAF Study 2**

Total RNA from differentiated human bronchial epithelial cells (HBECs) from controls was extracted and purified with a RNeasy Plus Micro Kit (Qiagen, Hilden, Germany). Samples with an RNA integrity number of greater than 9.0 were chosen for sequencing. Library preparation for RNA-seq was performed with the TruSeq Stranded mRNA Sample Prep Kit (Illumina, San Diego, Calif). Sequencing was performed on the Illumina HiSeq 4000.

**SIAF Study 3 (GSE112591)**

ILC1, ILC2 and ILC3 cell subpopulations from control individuals were obtained through a “2-round sorting” procedure, and processed as described in the original paper [2]. Briefly, 18 samples were analyzed using RNA sequencing on Illumina HiSeq 2500 platform.

**SIAF Study 4**

Skin tissue biopsies were collected from control individuals and atopic dermatitis patients (lesional and non-lesional sites in the upper arm/lower back area) as described in the original paper [3]. Briefly, RNA from obtained samples was extracted and purified and then sequenced (RNA-seq) on Illumina HiSeq 2500.

**SIAF Study 5 (GSE110551)**

Whole blood samples collected from healthy controls, asthma and COPD patients were stored at -80°C. BAL and bronchial biopsy samples from the same set of participants were harvested and frozen in RNALater solution. Next, samples were processed as described in the original paper [4]. Briefly, RNeasy Mini Kit was used to extract total RNA. Illumina HiSeq 2000 platform was used for sequencing of barcoded libraries with 51 bp paired-end reads for coverage of 50 million paired reads per sample.

**SIAF Study 6**

PBMCs were isolated from healthy infants (12-36 months of age) recruited from the AmaXhosa population in South Africa. RNA was isolated using RNeasy Plus Mini kit (Qiagen). Library was prepared with Illumina's TruSeq RNA stranded kit with polyA enrichment. RNA-Seq analysis was performed using HiSeq 4000.

**Study 7 (GSE107011)**

Immune cells were sorted and processed as described in the original paper [5]. Briefly, RNA aliquots were isolated. The cDNA libraries were prepared using modified SMARTSeq v2 protocol with the Illumina Nextera XT kit. The length distribution of the cDNA libraries was monitored using DNA High Sensitivity Reagent Kit (Perkin Elmer). RNA-Seq analysis was performed using Illumina HiSeq 2000.

**Study 8 (GSE134985)**

PBMCs from Tanzanian children were isolated and processed as described in the original paper [6]. RNA was isolated using RNeasy Plus Mini kit (Qiagen). Libraries were generated from samples with high RNA quality using TruSeq Stranded mRNA Library prep (Illumina) by Eurofins. Libraries were sequenced to an average depth of 24 million 1 × 50– base pair single-end reads per sample on HiSeq 2500 (Illumina, v4 chemistry).

**Study 9 (GSE79970)**

PBMCs from healthy adolescent were isolated and processed as described in the original paper [7]. Briefly, total RNA was extracted and purified. cDNA preparation was followed by RNAseq on Illumina HiSeq 2500.

**Study 10 (Stanford University)**

PBMCs from healthy adults were isolated. RNA was extracted, the libraries were sequenced on Illumina NextSeq500 instrument using 2x100bp paired-end reads, following the manufacturer's protocols.

**Study 11 (GSE114065)**

Children's naïve $CD4^+$ T cells were purified and processed as described in the original paper [8]. Briefly, total RNA was isolated from the purified cells. Library preparation was performed using the Illumina TruSeq Stranded mRNA Kit and libraries were sequenced on the Illumina HiSeq 4000 instrument.

**Data processing**

All RNA-seq data were processed with the same inhouse workflow available at https://github.com/uzh/ezRun. Significance threshold for differentially expressed genes was set to $p < 0.05$ and was calculated for the entire gene lists in each project. All calculations between different conditions were done using the edgeR R package. Spearman correlation coefficient was calculated using Hmisc R package, with the threshold for significance set to $\alpha = 0.05$. Correlation plots were done using Python's Seaborn library. Coexpression heatmaps as well as correlation heatmaps were done using the corrplot R package.

**Table S1. Summary of the ACE-2-, CD147- and CD26- related genes analyzed in the study**

| Pathway | Gene name | Protein symbol | Protein name |
|---|---|---|---|
| ACE2 | *ACE2* | ACE2 | Angiotensin-converting enzyme 2 |
| | *ACE* | ACE | Angiotensin-converting enzyme |
| | *TMPRSS2* | TMPRSS2 | Transmembrane protease serine 2 |
| | *SLC6A19* | S6A19, B(0)AT1 | Sodium-dependent neutral amino acid transporter B(0)AT1 |
| CD147 | *APH1A* | APH1A | Gamma-secretase subunit APH-1A |
| | *APH1B* | APH1B | Gamma-secretase subunit APH-1B |
| | *APOD* | APOD | Apolipoprotein D |
| | *BSG* | CD147, EMMPRIN | Basigin |
| | *CD44* | CD44 | CD44 |
| | *EGFR* | EGFR | Epidermal growth factor receptor |
| | *GP6* | GPVI | Platelet glycoprotein VI |
| | *ITGA3* | ITGA3 | Integrin alpha-3 |
| | *ITGA6* | ITGA6 | Integrin alpha-6 |
| | *ITGB1* | ITGB1, CD29 | Integrin beta-1 |
| | *JUP* | Plak | Junction plakoglobin |
| | *LGALS3* | Gal-3 | Galectin-3 |
| | *NCSTN* | Nicastrin | Nicastrin |
| | *NME1* | NDK A | Nucleoside diphosphate kinase A |
| | *NOD2* | NOD2 | Nucleotide-binding oligomerization domain-containing-2 |
| | *NXNL1* | NXNL1 | Nucleoredoxin-like protein 1 |
| | *PPIA* | CypA | Cyclophilin A |
| | *PPIB* | CypB | CYPB, Cyclophilin B |
| | *PSEN1* | PS-1 | Presenilin 1 |
| | *PSENEN* | PEN-2 | Gamma-secretase subunit PEN-2 |
| | *S100A9* | S100A9 | Protein S100A9 |
| | *SDC1* | SYND1 | Syndecan-1 |
| | *SLC16A1* | MCT1 | Monocarboxylate transporter 1 |

| | | | |
|---|---|---|---|
| CD147 | *SLC16A3* | MCT4 | Monocarboxylate transporter 4 |
| | *SLC16A4* | MCT5 | Monocarboxylate transporter 5 |
| | *SLC16A7* | MCT2 | Monocarboxylate transporter 2 |
| | *SLC16A8* | MCT3 | Monocarboxylate transporter 3 |
| | *SLC2A1* | GLUT1 | Solute carrier family 2, Facilitated glucose transporter member 1 |
| | *SLC3A2* | CD98 | 4F2 cell-surface antigen heavy chain |
| | *SLC7A5* | CD98 light chain | Large neutral amino acids transporter small subunit 1 |
| | *SPN* | CD43, GALGP | Leukosialin |
| NFATs | *NFAT5* | NF-AT5 | Nuclear factor of activated T-cells 5 |
| | *NFATC1* | NF-ATc1 | Nuclear factor of activated T-cells, cytoplasmic 1 |
| | *NFATC2* | NF-ATc2 | Nuclear factor of activated T-cells, cytoplasmic 2 |
| | *NFATC3* | NF-ATc3 | Nuclear factor of activated T-cells, cytoplasmic 3 |
| | *NFATC4* | NF-ATc4 | Nuclear factor of activated T-cells, cytoplasmic 4 |
| CD26 | *DPP4* | CD26 | Dipeptyl peptidase 4 |

**Table S2**. **Summary of purity, markers and methods of purification in immune cell analysis**

| Cell type | Purity | Markers | Methods of purification | Reference |
|---|---|---|---|---|
| Neutrophils | >98% | CD3-CD19-CD45+SCC-A+(high) CD16+ | PBMCs were firstly separated into CD3+ and CD3- populations using magnetic beads. The CD3+ fraction was then split for different T cell staining while the CD3- fraction was split for B cell and progenitor-cell staining, or for monocyte, DCs, NK cells and LD granulocyte staining. After staining, the immune cells were sorted using a BD Influx, a FACS Aria 5 or a FACS Aria 4. | [5] |
| Classical monocytes | | CD3-CD19-CD45+CD11c+CD14+CD16- | | |
| Plasmacytoid dendrite cells | | CD3-CD19-CD45+HLA-DR+CD123+ | | |
| Natural killer cells | | CD3-CD19-CD45+CD16+CD56+ | | |
| Naïve $CD4^+$ T cells | | CD3+CD4+CCR7+CD45RA+ | | |
| Terminal effector $CD4^+$ T cells | | CD3+CD4+CCR7+CD45RA- | | |
| Naïve $CD8^+$ T cells | | CD3+CD8+CCR7+CD45RA+ | | |
| Effector memory $CD8^+$ T cells | | CD3+CD8+CCR7-CD45RA- | | |
| Naïve B cells | | CD3-CD56-CD16-CD14-CD45+CD19+CD27-IgD+ | | |
| Plasmablasts | | CD3-CD56-CD16-CD14-CD45+CD19+CD27+IgD-CD38+(high) | | |
| ILC1 | 86%-96% | CD45+Lin-CD127+CD161+CRTH2-c-Kit- | Robust “2-round sorting” procedure by using flow cytometry with modified Mjösberg protocol | [2] |
| ILC2 | | CD45+Lin-CD127+CD161+CRTH2+c-Kit- | | |
| ILC3 | | CD45+Lin-CD127+CD161+CRTH2-c-Kit+ | | |

**Table S3. Summary of projects and databases analyzed in the manuscript**

<table>
<tr><th>Database</th><th>Cell Type</th><th>Condition</th><th>GEO</th><th colspan="2">Number of subjects</th><th>Age</th><th>Reference</th></tr>
<tr><td>SIAF 1</td><td>HBECs</td><td>(in vitro)</td><td>n/a</td><td colspan="2">Control = 5<br>Asthma = 6<br>COPD = 5</td><td>adults</td><td>[1]</td></tr>
<tr><td>SIAF 2</td><td>HBECs</td><td>(in vitro)</td><td>n/a</td><td colspan="2">Control = 5</td><td>adults</td><td>n/a</td></tr>
<tr><td>SIAF 3</td><td>ILC1, ILC2, ILC3</td><td>(ex vivo)</td><td>GSE112591</td><td colspan="2">ILC1 = 6<br>ILC2 = 6<br>ILC3 = 6</td><td>adults</td><td>[2]</td></tr>
<tr><td>SIAF 4</td><td>Skin biopsy</td><td>(ex vivo)</td><td>n/a</td><td colspan="2">Control = 6<br>Atopic dermatitis (non lesional) = 11<br>Atopic dermatitis (lesional) = 11</td><td>adults</td><td>[3]</td></tr>
<tr><td rowspan="3">SIAF 5</td><td>Bronchial biopsy</td><td rowspan="3">(ex vivo)</td><td rowspan="3">GSE110551</td><td>Control = 16<br>Asthma = 22<br>COPD = 3<br>Non-obese = 20<br>Obese = 21</td><td>Normotension = 32<br>Hypertension = 9<br>Non-smoker = 19<br>Smoker = 21<br>Female = 14<br>Male = 27</td><td rowspan="3">20-60 years</td><td rowspan="3">[4]</td></tr>
<tr><td>BAL</td><td>Control = 16<br>Asthma = 22<br>COPD = 2<br>Non-obese = 19<br>Obese = 21</td><td>Normotension = 31<br>Hypertension = 9<br>Non-smoker=19<br>Smoker = 20<br>Female = 14<br>Male = 26</td></tr>
<tr><td>Blood</td><td>Control = 17<br>Asthma = 21<br>COPD = 3<br>Non-obese = 20<br>Obese = 21</td><td>Normotension = 32<br>Hypertension = 9<br>Non-smoker = 19<br>Smoker = 21<br>Female = 14<br>Male = 27</td></tr>
<tr><td>SIAF 6</td><td>Children PBMCs</td><td>(ex vivo)</td><td>n/a</td><td colspan="2">Control children = 14</td><td>12-36 months</td><td>n/a</td></tr>
<tr><td>Study 7</td><td>neutrophils, classical monocytes, plasmacytoid dendrite cells, natural killer cells, naïve CD4+ T cells, terminal effector CD4+ T cells, naïve CD8+ T cells, effector memory CD8+ T cells,</td><td>(ex vivo)</td><td>GSE107011</td><td colspan="2">Neutrophils = 4<br>Classical monocytes = 4<br>Plasmacytoid dendrite cells = 4<br>Natural killer cells = 4<br>Naïve CD4+ T cells = 4<br>Terminal effector CD4+ T cells = 2<br>Naïve CD8+ T cells = 4<br>Effector memory CD8+ T cells = 4<br>Naïve B cells = 4<br>Plasmablasts = 4</td><td>20-35 years</td><td>[5]</td></tr>
</table>

| | | | | | | |
|---|---|---|---|---|---|---|
| | naïve B cells, plasmablasts | | | | | |
| Study 8 | Children PBMCs | *(ex vivo)* | GSE134985 | Control children = 21 | 5-17 months | [6] |
| Study 9 | Adolescent PBMCs | *(ex vivo)* | GSE79970 | Control adolescent = 16 | 4-16 years | [7] |
| Study 10 | Adults PBMCs | *(ex vivo)* | n/a | Control adult = 19 | 16-67 years | n/a |
| Study 11 | Children naïve CD4+ T cells | *(ex vivo)* | GSE114065 | Control children = 18 | 12 months | [8] |

### Online References


1. Wang, M.; Tan, G.; Eljaszewicz, A.; Meng, Y.; Wawrzyniak, P.; Acharya, S.; Altunbulakli, C.; Westermann, P.; Dreher, A.; Yan, L., et al. Laundry detergents and detergent residue after rinsing directly disrupt tight junction barrier integrity in human bronchial epithelial cells. *J Allergy Clin Immunol* **2019**, *143*, 1892-1903, doi:10.1016/j.jaci.2018.11.016.
2. Li, S.; Morita, H.; Sokolowska, M.; Tan, G.; Boonpiyathad, T.; Opitz, L.; Orimo, K.; Archer, S.K.; Jansen, K.; Tang, M.L.K., et al. Gene expression signatures of circulating human type 1, 2, and 3 innate lymphoid cells. *J Allergy Clin Immunol* **2019**, *143*, 2321-2325, doi:10.1016/j.jaci.2019.01.047.
3. Altunbulakli, C.; Reiger, M.; Neumann, A.U.; Garzorz-Stark, N.; Fleming, M.; Huelpuesch, C.; Castro-Giner, F.; Eyerich, K.; Akdis, C.A.; Traidl-Hoffmann, C. Relations between epidermal barrier dysregulation and Staphylococcus species-dominated microbiome dysbiosis in patients with atopic dermatitis. *J Allergy Clin Immunol* **2018**, *142*, 1643-1647 e1612, doi:10.1016/j.jaci.2018.07.005.
4. Michalovich, D.; Rodriguez-Perez, N.; Smolinska, S.; Pirozynski, M.; Mayhew, D.; Uddin, S.; Van Horn, S.; Sokolowska, M.; Altunbulakli, C.; Eljaszewicz, A., et al. Obesity and disease severity magnify disturbed microbiome-immune interactions in asthma patients. *Nat Commun* **2019**, *10*, 5711, doi:10.1038/s41467-019-13751-9.
5. Monaco, G.; Lee, B.; Xu, W.; Mustafah, S.; Hwang, Y.Y.; Carre, C.; Burdin, N.; Visan, L.; Ceccarelli, M.; Poidinger, M., et al. RNA-Seq Signatures Normalized by mRNA Abundance Allow Absolute Deconvolution of Human Immune Cell Types. *Cell Rep* **2019**, *26*, 1627-1640.e1627, doi:10.1016/j.celrep.2019.01.041.
6. Hill, D.L.; Carr, E.J.; Rutishauser, T.; Moncunill, G.; Campo, J.J.; Innocentin, S.; Mpina, M.; Nhabomba, A.; Tumbo, A.; Jairoce, C., et al. Immune system development varies according to age, location, and anemia in African children. *Sci Transl Med* **2020**, *12*, doi:10.1126/scitranslmed.aaw9522.
7. Wong, L.; Jiang, K.; Chen, Y.; Hennon, T.; Holmes, L.; Wallace, C.A.; Jarvis, J.N. Limits of Peripheral Blood Mononuclear Cells for Gene Expression-Based Biomarkers in Juvenile Idiopathic Arthritis. *Sci Rep* **2016**, *6*, 29477, doi:10.1038/srep29477.
8. Martino, D.; Neeland, M.; Dang, T.; Cobb, J.; Ellis, J.; Barnett, A.; Tang, M.; Vuillermin, P.; Allen, K.; Saffery, R. Epigenetic dysregulation of naive CD4+ T-cell activation genes in childhood food allergy. *Nat Commun* **2018**, *9*, 3308, doi:10.1038/s41467-018-05608-4.
